# Structured and disordered regions of Ataxin-2 contribute differently to the specificity and efficiency of mRNP granule formation

**DOI:** 10.1101/2022.02.15.480566

**Authors:** Arnas Petrauskas, Daniel L. Fortunati, Amanjot Singh, Arvind Reddy Kandi, Sai Shruti Pothapragada, Khushboo Agrawal, Joern Huelsmeier, Jens Hillebrand, Georgia Brown, Dhananjay Chaturvedi, Jongbo Lee, Chunghun Lim, Georg Auburger, K. VijayRaghavan, Mani Ramaswami, Baskar Bakthavachalu

## Abstract

Ataxin-2 (*ATXN2*) is a gene implicated in spinocerebellar ataxia type II (SCA2), amyotrophic lateral sclerosis (ALS) and Parkinsonism. The encoded protein is a therapeutic target for ALS and related conditions. ATXN2 (or Atx2 in insects) can function in translational activation, translational repression, mRNA stability and in the assembly of mRNP-granules, a process mediated by intrinsically disordered regions (IDRs). Previous work has shown that the LSm (Like-Sm) domain of Atx2, which can help stimulate mRNA translation, antagonizes mRNP-granule assembly. Here we advance these findings through a series of experiments on *Drosophila* and human Ataxin-2 proteins. Results of Targets of RNA-Binding Proteins Identified by Editing (TRIBE), co-localization and immunoprecipitation experiments indicate that a polyA-binding protein (PABP) interacting, PAM2 motif of Ataxin-2 may be a major determinant of the mRNA and protein content of Ataxin-2 mRNP granules. Transgenic experiments in *Drosophila* indicate that while the Atx2-LSm domain may protect against neurodegeneration, structured PAM2- and unstructured IDR- interactions both support Atx2-induced cytotoxicity. Taken together, the data lead to a proposal for how Ataxin-2 interactions are remodelled during translational control and how structured and non-structured interactions contribute differently to the specificity and efficiency of RNP granule condensation as well as to neurodegeneration.

## INTRODUCTION

mRNP granules are intriguing, dynamic membrane-less organelles containing translationally repressed mRNAs, RNA-binding proteins (RBPs), molecular chaperones and a variety of other cellular proteins (Buchan, 2014; Formicola *et al*, 2019; Kiebler & Bassell, 2006; Knowles *et al*, 1996; Martin & Ephrussi, 2009). The formation and composition of mRNP assemblies are determined by base-pairing interactions between mRNAs, protein-protein interactions and RBP-RNA interactions, whose respective contributions may vary across granule types and physiological states (Bevilacqua *et al*, 2022; Matheny *et al*, 2021; Van Treeck & Parker, 2018; Van Treeck *et al*, 2018). Stress granules (SGs) are particularly well-studied granules that form when cellular stress signals mediated by eIF2α kinase activation (Kedersha *et al*, 1999) cause individual mRNPs to arrest in translation and condense into multi-mRNP assemblies (Ivanov *et al*, 2019; Kedersha & Anderson, 2007; Youn *et al*, 2019). Mutations in mRNP granule proteins, including TDP-43, FUS, Ataxin-2, hnRNPA1, hnRNPA2B1, EWSR1, have been associated with ALS and/or other forms of neurodegenerative disease (Cirulli *et al*, 2015; Couthouis *et al*, 2012; Elden *et al*, 2010; Ginsberg *et al*, 1998; Kim *et al*, 2013; Liu *et al*, 2017; Taylor *et al*, 2016; Wolozin & Ivanov, 2019). For this reason, and because TDP-43 and other stress-granule protein aggregates are components of protein inclusions found in ALS and Frontotemporal dementia (FTD), the regulation and cellular functions of stress granules have been topics of considerable fundamental and clinical interest (Cao *et al*, 2020; Li *et al*, 2013; Mallucci *et al*, 2020; Protter & Parker, 2016; Wang *et al*, 2020; Wheeler *et al*, 2016; Wolozin & Ivanov, 2019).

The cast of intermolecular interactions required for mRNP-granule assembly and the precise sequence with which they occur are not yet elucidated (Khong & Parker, 2020; Van Treeck & Parker, 2018). However, many studies show that intrinsically disordered regions (IDRs) found on mRNP-granule proteins contribute substantially to RNP granule assembly (Andrusiak *et al*, 2019; Ash *et al*, 2021; Calabretta & Richard, 2015; Decker *et al*, 2007; Gilks *et al*, 2004; Järvelin *et al*, 2016; Kim *et al*, 2021; Yang *et al*, 2020). In biochemical experiments, such IDRs show the ability to phase separate into liquid-like assemblies (Babinchak & Surewicz, 2020; Han *et al*, 2012; Hyman *et al*, 2014; Kato *et al*, 2012; Lin *et al*, 2017; Murray *et al*, 2017; Murthy *et al*, 2019; Shin & Brangwynne, 2017; Strome & Wood, 1982; Toretsky & Wright, 2014; Yang *et al*., 2020) The accessibility or activities of IDRs can be tightly regulated by posttranslational modifications, allowing rapid physiological and spatial control over granule assembly and disassembly (Ash *et al*., 2021; Bah & Forman-Kay, 2016; Bah *et al*, 2015; Berlow *et al*, 2015; Hofweber & Dormann, 2019; Kwon *et al*, 2013; Rayman *et al*, 2018; Saito *et al*, 2019; Yang *et al*., 2020).

An important observation is that most IDRs also have the ability to transition from liquid-like states into solid, beta-sheet rich, amyloid-fibrils *in vitro*, particularly at high concentrations achieved in the liquid-phase (Alberti *et al*, 2019; Li *et al*., 2013; Murray *et al*., 2017; Patel *et al*, 2015; Ramaswami *et al*, 2013). This, and studies showing that inhibitors of eIF2α kinase or downstream events including SG formation can be protective in animal models of neurodegenerative disease (Chou *et al*, 2017; Halliday *et al*, 2017; Sidrauski *et al*, 2015; Wong *et al*, 2018; Zyryanova *et al*, 2021) have led to a conceptual framework in which: (a) mRNP granules provide a microenvironment where pathogenic protein seeds can form and grow (Bakthavachalu *et al*, 2018; Mandrioli *et al*, 2020; Patel *et al*., 2015); (b) increased misfolded-protein loads result in inclusion formation, chronic stress signalling and reduced protein translation (Hetz *et al*, 2020; Preissler & Ron, 2019); (c) increased demand on protein handling systems results in multiple cellular defects, notably in the functions of membrane-less organelles (Alberti *et al*, 2017; Azkanaz *et al*, 2019; Jiang *et al*, 2020; Latonen, 2019; Schuller *et al*, 2021). In particular, aberrant SG formation also results in nuclear transport defects which may contribute to cell death and toxicity (Hochberg-Laufer *et al*, 2019; Zhang *et al*, 2018).

Particularly strong support for the role of RNP granule formation in promoting disease comes from studies of Ataxin-2. Loss of Ataxin-2 is cytoprotective in yeast TDP-43 and *Drosophila* TDP-43 or C9ORF72 or Tau models of cytotoxicity (Bakthavachalu *et al*., 2018; Becker *et al*, 2017; Elden *et al*., 2010; Huelsmeier *et al*, 2021; Kim *et al*, 2014; Lee *et al*, 2016; Shulman & Feany, 2003). In mouse models for SCA2 or ALS, either genetic loss of *ATXN2* or delivery of antisense oligonucleotides (ASOs) targeting *ATXN2* in the central nervous system, reduced aggregation of TDP-43, increased animal survival and improved motor function (Becker *et al*., 2017; Scoles *et al*, 2017). These observations have led to ASOs against human *ATXN2* being developed and approved for clinical trials (Biogen, 2021).

Given Ataxin-2’s therapeutic significance and multiple roles in biology, it is particularly important to determine which molecular activities of the protein are relevant to disease and to its various biological functions (Kim *et al*, 2020). Across species, Ataxin-2 has three conserved regions: a Like-Sm (LSm) domain, an LSm-associated domain (LSm-AD) and a PAM2 motif, which is flanked by extended intrinsically disordered regions (IDRs) (Boeynaems *et al*, 2021). Detailed work in *Drosophila* has shown that a c-terminal IDR of Atx2 is selectively required for mRNP assembly into granules (Bakthavachalu *et al*., 2018). Parallel experiments showing that the IDR is also required for cytotoxicity in *Drosophila* FUS, C9ORF72 and Huntington’s disease models suggest RNP-granule formation to be a significant mechanism by which Atx2 promotes neurodegeneration (Bakthavachalu *et al*., 2018; Huelsmeier *et al*., 2021). A recent discovery that the Atx2-LSm domain antagonizes IDR-function has led to a model in which the Atx2 cIDR: (a) does not support mRNP assembly when Atx2 is associated with actively translating mRNAs through Atx2-LSm domain interactions; (b) becomes accessible and active in mediating mRNP assembly when LSm-domain interactions break and mRNA translation stalls (Boeynaems *et al*., 2021; Singh *et al*, 2021).

We now present a series of experiments further elaborating mechanisms by Ataxin-2 functions in mRNA translation and mRNP assembly. These show that the PAM2 motif of Ataxin-2 and its interactions with PABP are not essential for granule assembly but are required to efficiently recruit Atx2-target mRNAs and specific protein components into Ataxin-2 granules. When taken together with other findings (Boeynaems *et al*., 2021; Kim *et al*., 2014; Satterfield & Pallanck, 2006; Singh *et al*., 2021), our observations indicate that PAM2 binding to PABP on the polyA tail of mRNAs helps specify the composition of Ataxin-2 granules. We propose an early role for PAM2:PABP interactions working in coordination with the LSm domain to support mRNA translation and thereby oppose the mRNP formation (Boeynaems *et al*., 2021); as well as a later role in escorting translationally-stalled PAM2:PABP linked mRNAs into mRNP granules. *In vivo* experiments analysing motor decline in transgenic *Drosophila* indicate that the PAM2:PABP interactions also support the progression of the neurodegenerative process. We provide new evidence for fresh insight into the enigmatic role of mRNP assembly in neurodegeneration.

## RESULTS

### The structured PAM2 domain of Atx2 is necessary for the correct mRNA and protein content of Atx2 granules

A recent eLife publication used Targets of RNA-Binding Proteins Identified by Editing (TRIBE) technology to identify mRNAs associated with Atx2 in the *Drosophila* adult brain (Singh *et al*., 2021). *In vivo*, the ability of an Atx2-fusion with ADARcd (the catalytic domain of an RNA-editing enzyme, ADAR), to edit a group of 256 target mRNAs was found to be dependent on the presence of the Atx2-cIDR, previously shown to be necessary for the formation of neuronal mRNP granules *in vivo*. In contrast, Atx2-ADARcd mutants lacking the LSm domain, both edited Atx2 TRIBE target RNAs and formed mRNP granules in cultured *Drosophila* S2 cells more efficiently than the wild-type. Thus, Atx2-ADARcd editing of target mRNAs occurs in and is reflective of mRNP granule assembly. While demonstrating a role for LSm-domain interactions in antagonizing cIDR mediated granule assembly, these observations did not address mechanisms by which Atx2 target mRNAs are selected, or whether and how Atx2 played any role in determining the composition of RNP granules. New experiments presented here address these outstanding questions.

Previous TRIBE analyses showed that LSm and LSm-AD regions have no major role in the recognition or selection of the Atx2-target mRNAs (Singh *et al*., 2021). We therefore tested whether the third conserved region of Ataxin-2, a PAM2 motif known to associate with PABP (polyA binding protein), played any role in this process (Jiménez-López & Guzmán, 2014; Kaehler *et al*, 2012).

We used Gal80^ts^-controlled *elav-Gal4* to express Atx2ΔPAM2-ADARcd (deleted for the PAM2 motif) in brains of adult *Drosophila* for 5 days and used RNA-Seq to identify edited RNAs in polyA selected brain mRNA and compare it with Atx2-ADARcd using procedures described earlier (Figure 1A) (McMahon *et al*, 2016). ADAR-edits, which converts Adenosine to Inosine on RNAs, are identified as A to G changes in TRIBE analyses. Each sample was sequenced to obtain 20 million reads (Supplementary table 1). The edits were only considered from the regions of the transcriptome that contained at least 20 reads. Genes with edits identified at a threshold above 15% in two biological replicates were considered as high-confidence true targets. We compared edit frequency and edited-gene identity in the brains of flies expressing Atx2ΔPAM2-ADARcd with those in brains expressing Atx2-ADARcd.

**Figure 1:**
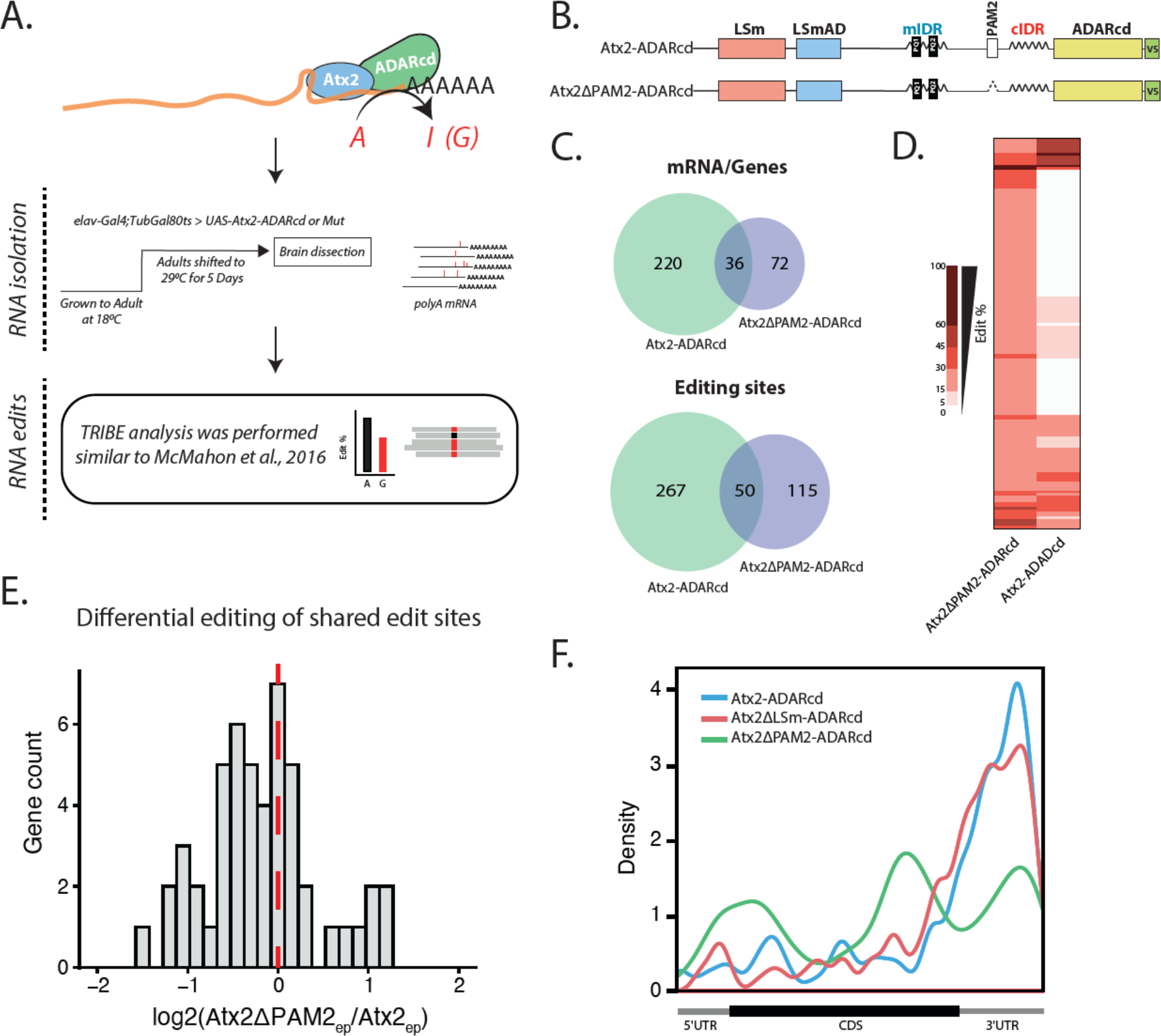
The PAM2 domain facilitates the selection of Atx2 RNA targets. (A). Flowchart depicting the TRIBE analyses pipeline. Atx2ΔPAM2-ADARcd was expressed in adult *Drosophila* brains. Total brain RNA was isolated and RNA edits were identified and compared to Atx2-ADARcd, similar to Singh et al 2021. (B) Domain map of Atx2-ADARcd constructs used for TRIBE analysis. (C) Comparisons of genes and edits identified by TRIBE between Atx2-ADARcd and Atx2ΔPAM2-ADARcd targets. (D) Most Atx2ΔPAM2 targets identified by TRIBE are unique and not edited in Atx2WT, suggesting that these new targets bound by Atx2ΔPAM2 are not native Ataxin-2 granule targets. (E) Comparisons of the editing efficiency ratio of common edits between Atx2WT vs Atx2ΔPAM2 show a much lower editing efficiency in Atx2ΔPAM2 compared to Atx2WT. (F) PAM2 deletion results in loss of 3’UTR specificity seen in Atx2WT and LSm deletion TRIBE target mRNAs. Atx2WT and Atx2ΔLSm-ADARcd data are extracted from (Singh et al 2021).

In contrast to Atx-2 forms lacking LSm or LSm-AD domains (Singh *et al*., 2021), Atx2ΔPAM2-ADARcd edited significantly fewer RNA targets than wild-type Atx2-ADARcd (108 genes and 165 edits vs 256 genes and 317 edits, Figure 1B, C and Supplementary table 2). More striking, the cohort of mRNAs edited by the ΔPAM2 mutant form differed extensively from the largely overlapping cohorts edited by either wild-type forms of Atx2 (Figure 1C). Of the 108 genes edited by Atx2ΔPAM2-ADARcd, 36 were also targets of Atx2-ADARcd, the remaining 72 were unique. (Figure 1C, D and Supplementary table 2). 50 edit sites were common between the Atx2ΔPAM2 and Atx2WT targets. Those sites were edited with much lower efficiency in Atx2ΔPAM2 as compared to Atx2WT (Figure 1E).

The location of edits made by Atx2ΔPAM2-ADARcd also differed dramatically as to where they occurred relative to the coding sequences of the target mRNAs (Figure 1F). While edits made by wild-type and ΔLSm forms of Atx2-ADARcd were greatly enriched in the 3’UTR of the mRNAs, Atx2ΔPAM2 targets were edited indiscriminately all along the mRNA length (Figure 1F).

Taken together, these data identify the PAM2 motif as necessary for Atx2 engagement with its correct mRNA targets. The PAM2 motif interacts with PABP, which binds polyA tracts at the 3’ end of mRNAs (Deo *et al*, 1999). Therefore, the data point to a role for the structured PAM2:PABP interaction in guiding the association of Atx2 with mRNAs and for subsequent inclusion of these mRNAs in Atx2-containing granules.

If Atx2-ADARcd edits of target mRNAs occur predominantly in the mRNP granules (Singh *et al*., 2021), then the ability of Atx2ΔPAM2-ADARcd to edit some target mRNAs would suggest that the PAM2 motif is not essential for mRNP granule formation *per se.* To examine this, we expressed wild-type and ΔPAM2 mutant forms of GFP-tagged Atx2 under control of the native genomic promoter in *Drosophila* S2R+ cells. Atx2 overexpression in S2 cells induced the formation of mRNP granules closely related to SGs, containing endogenous Atx2 and various SG proteins as previously reported (Figure 2A) (Bakthavachalu *et al*., 2018; Singh *et al*., 2021). Similar expression of Atx2ΔPAM2-GFP also induced granule formation. However, these granules were compositionally distinct from those induced by Atx2-GFP. While they clearly contained some SG markers present on Atx2-granules, e.g., Me31B and Rox8 (*Drosophila* homologs of DDX6 and TIA1), they did not contain others such as PABP, Caprin and dFMRP (Figure 2B).

**Figure 2:**
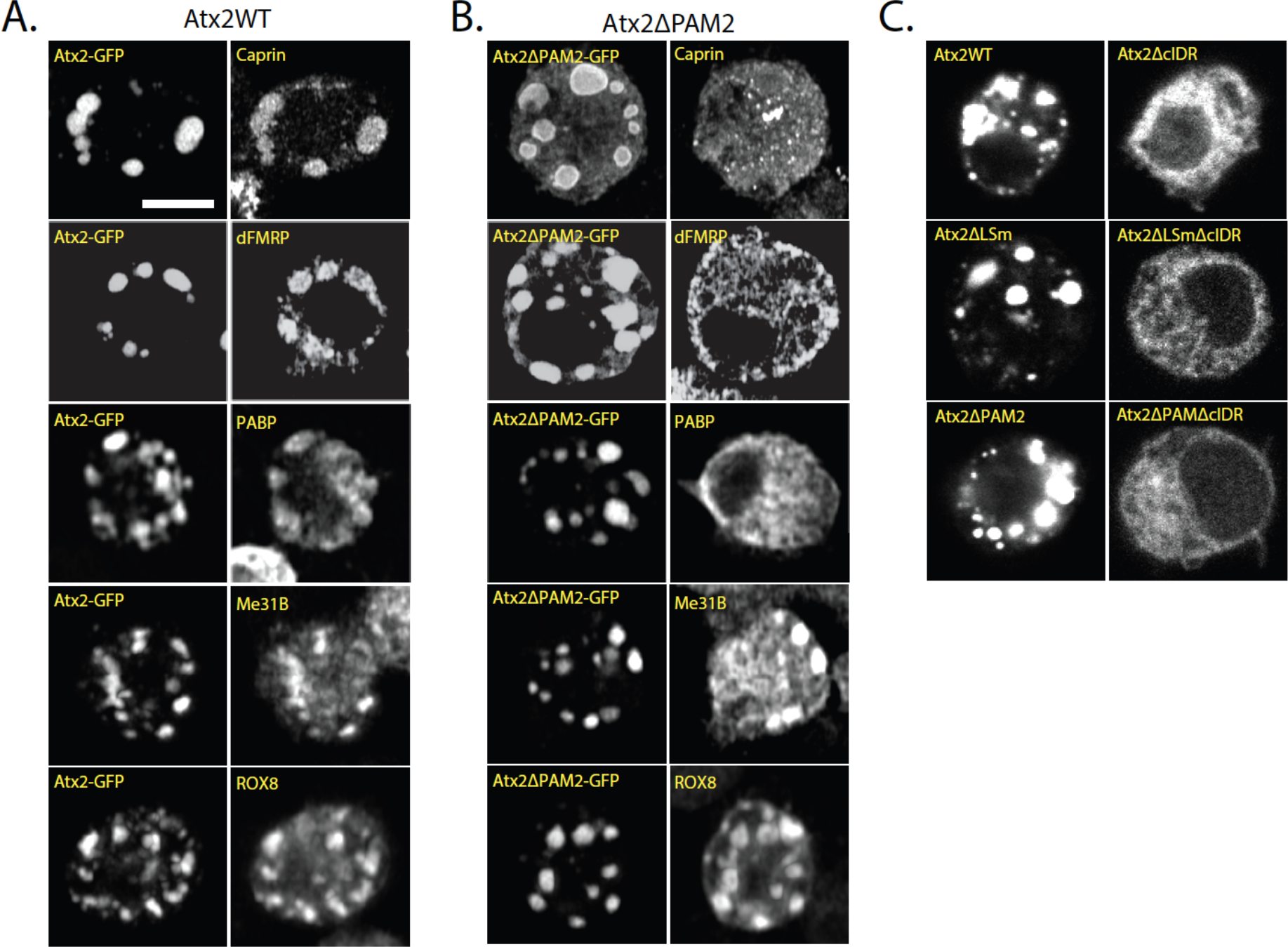
Presence of the PAM2 domain affects the protein composition of Atx2-GFP granules in S2 cells. (A) Over-expression of Atx2-GFP in unstressed *Drosophila* S2 cells induces the formation of Atx2-GFP granules to which various SG markers co-localize. (B) Deletion of the PAM2 affects the Atx2-GFP granule composition. Over-expression of Atx2ΔPAM2-GFP in S2 cells still induces the formation of granules, but some SG markers fail to co-localize in these, notably dFMR, Caprin and PABP. (C) Atx2-GFP granule formation in S2 cells relies primarily on the cIDR. Deletion of the cIDR in Atx2WT, Atx2ΔPAM2 and Atx2ΔLSm, removes their ability to form granules upon overexpression. See Supplemental Figure 1, A-B, for quantification. The scale bar in (A) applies to (B) and (C). Scale bar = 5 μm,

RNP-granules induced by expression of wild-type, LSm and PAM2 deficient forms of Atx2-GFP required the presence of the c-terminal IDR (Figure 2C). Thus, while largely dispensable for efficient mRNP assembly, the PAM2 domain plays a significant role in determining both mRNA and protein components of mRNP granules. One possibility is that the PAM2 motif directly recruits PABP and associated mRNAs to granules and indirectly recruits other proteins through their interactions with either PABP or mRNAs brought to RNP granules through Atx2-PAM2:PABP interactions.

### PAM2:PABP interactions are sufficient for Atx2 to associate with stress granules

We wanted to directly confirm Ataxin-2 PAM2 motif interactions with PABP and analyse their relevance to RNP granule assembly. For this, we generated constructs encoding SNAP-epitope tagged variants of Atx2. These were radically truncated forms of *Drosophila* and human Ataxin-2 proteins containing only the LSm, LSm-AD and PAM2 elements and lacking all unstructured regions of the protein. The structured elements are connected via flexible linkers (Figure 3A). These “Mini-Ataxin-2” constructs and their domain-deleted forms allowed us to separate functions of the structured regions of Ataxin-2 from those of the remaining extended disordered regions. A similar approach has been previously shown for MeCP2 (Tillotson *et al*, 2017). We identified key residues involved in *Drosophila* Atx2-PAM2:PABP interactions based on a previously solved crystal structure of a strongly conserved mammalian PAM2:PABPC1-MLLE domain complex (Kozlov *et al*, 2010; Xie *et al*, 2014) (Figure 3B). Residues leucine 914 and phenylalanine 921 (L914 and F921) in the human ATXN2-PAM2 motif are predicted to contact the PABPC-MLLE domain and of these, F921 has been shown to be required for the PABPC-ATXN2 interaction (Inagaki *et al*, 2020). These residues (L859 and F866, respectively) are perfectly conserved in fly Atx2 (Supplementary Figure 3). In order to allow more precise disruption of PAM2:PABP interactions and avoid potential unknown secondary effects of larger PAM2 motif deletions, we additionally generated mini Ataxin-2 constructs where these PABP-contacting residues were singly or doubly altered to alanine. We used these constructs for co-immunoprecipitation (Figure 3) and co-localization (Figure 4) analyses to examine the contribution of PAM2:PABP/PABPC1 interactions in RNP-granule formation.

**Figure 3:**
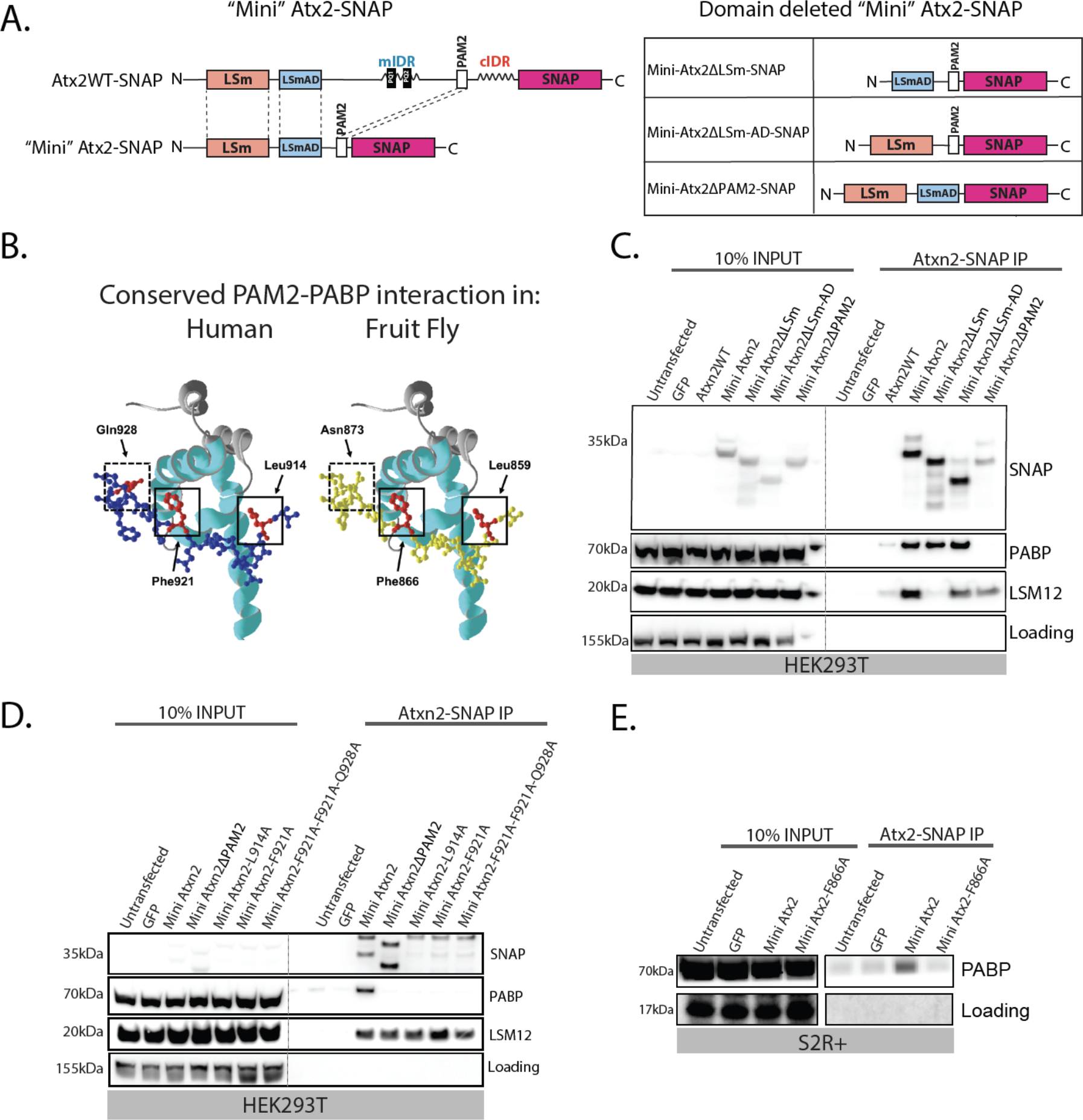
A minimized Ataxin-2 construct containing only the known structured domains maintains the ability to interact with PABP and LSM12. (A) Schematic of SNAP-tagged full length, minimal, and minimal domain-deleted constructs of fly Atx2 used to isolate the function of the structured domains of the protein without interference from IDR-mediated interactions. Human and *Drosophila* Ataxin-2 LSm, LSm-AD and PAM2 domains show high amino acid sequence similarity (Clustal Ω) of 70%, 82% and 87% respectively. This suggests conserved and specific function of these structured domains. (B) Structural model of the PABP MLLE domain (ribbon) showing the near-perfect structural similarity of the human ATXN2 PAM2 domain (blue, uniprot ID: Q99700) with the *Drosophila* Atx2 PAM2 domain (yellow, uniprot ID: Q8SWR8). The key interacting residues are highlighted in red. (C) Human minimized ATXN2 SNAP IP-WB from HEK293T cells probing for PABP and LSM12 showing the effects of different domain deletions. The PAM2 domain is necessary and sufficient for the ATXN2-PABP interaction, while the LSm domain is necessary and sufficient for the ATXN2-LSM12 interaction. (D) Point-mutations targeting key interacting residues of the PAM2 domain were predicted to replicate the effect of a full PAM2 deletion in the minimized Atx2 construct. Human minimized ATXN2-SNAP IP WB from HEK293T cells showing that mutating either of the key hydrophobic residues L914 or F921 in the PAM2 domain is sufficient to prevent its interaction with PABP. The interaction with LSM12 is unaffected by the point mutations. (E) *Drosophila* minimized Atx2-SNAP IP-WB from S2 cells. An analogous PAM2 domain point mutation on F866 blocks the Atx2-PABP interaction.

**Figure 4:**
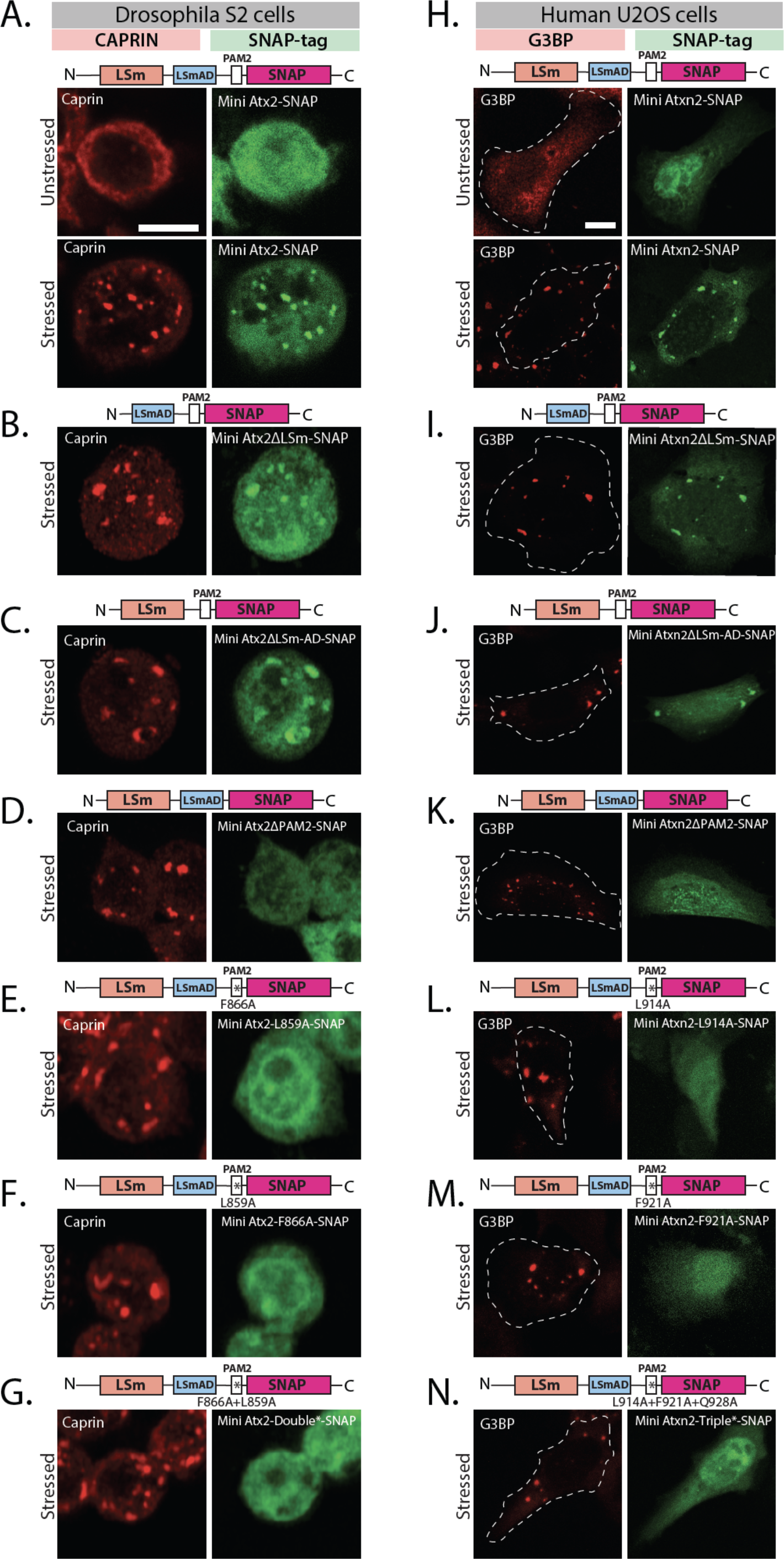
The structured PAM2 domain is necessary and sufficient for Ataxin-2 recruitment to Stress Granules in both *Drosophila* and human cells. (A) In *Drosophila* S2 cells mini-Atx2-SNAP (green) is recruited to SGs induced by arsenite. (B) Deletion of LSm or (C) LSm-AD domains has no significant effect in the arsenite-induced SG recruitment. (D) The presence of PAM2 domain, and specifically its key PABP-interacting amino acids (E-G), is necessary for Atx2 recruitment to SGs. Caprin (red) was used as SG granule marker, scale bar = 5 μm, (H) Human mini-ATXN2-SNAP (green) is recruited to arsenite-induced SGs in human U2OS cells. (I) Deletion of LSm or (J) LSm-AD domains has no effect on the arsenite-induced SG recruitment of ATXN2. (K) Deletion of the PAM2 domain, and specifically its PABP-interacting amino acids (L-N), are necessary for ATXN2 recruitment to SGs. G3BP1 (red) was used as SG marker; scale bar = 10 μm. Schematics above images indicate the domain deletions or amino acid mutations that were made in the different Ataxin-2 constructs. See SSupplemental Figure 1C for quantification.

We expressed SNAP-tagged wild-type and mutant forms of mammalian and *Drosophila* mini-Ataxin-2 in HEK293T and S2 cells, respectively. We tested which Ataxin-2 domains were required for SNAP substrate beads to successfully immunoprecipitate Ataxin-2 complexes containing PABPC1/PABP from cell lysates (Figure 3C-E). Both LSM12 and PABPC1 proteins were co-immunoprecipitated with mammalian mini-ATXN2. However, PABPC1 co-immunoprecipitation was selectively lost when the PAM2 domain was deleted or if predicted PABP-contact residues in the PAM2 domain were mutated (Figure 3C). Similar to the human homolog, fly mini-Atx2-SNAP also required the presence of its PAM2 motif with both predicted contact residues intact for immunoprecipitation of PABP from *Drosophila* S2 cell lysates (Figure 3D). Taken together with previous observations, these data support a potential sequence of molecular events. In unstressed cells, PAM2 domain interaction with PABP help position Ataxin-2 at the 3’-end of mRNAs while LSM-domain association with LSM12 stimulate translation of these mRNAs; under stress conditions (or Ataxin-2 overexpression), translation is arrested and the cIDR domain freed to mediate interactions that facilitate the formation of condensed RNP granules (see Discussion)

Single mRNAs usually associate with multiple PABP molecules because their polyA tails are considerably longer than the ∼24 bases required for PABP binding (Mangus *et al*, 2003). Thus, in cells expressing endogenous and mini Ataxin-2, mRNAs could have both forms associated with their polyA tails. In response to stress, mini Ataxin-2 would be expected to move into SGs whose formation is facilitated by IDRs on endogenous Ataxin-2 associated with the common target RNAs. We examined this possibility in cells before and after oxidative stress.

*Drosophila* and human mini-Ataxin-2-SNAP, expressed in fly S2 or human U2OS cells respectively, were diffusely localized in the cytoplasm and neither induced formation of Ataxin-2 foci. However, when cells were exposed to sodium arsenite to induce SG formation, SNAP-tagged mini-Atx2 (Figure 4A) and mini-ATXN2 (Figure 4H) were robustly recruited to stress granules. Thus, association of Ataxin-2 with mRNP-granule components may be achieved by structured domain interactions alone, independently of IDRs required for mRNP assembly into granules.

Further experiments examined which of the LSm, LSm-AD and/or PAM2 domains were necessary for mini-Ataxin-2 to associate with stress granules. Mammalian and *Drosophila* mini-Ataxin-2 forms missing the LSm or LSm-AD domains could still be found in stress granules (Figure 4B-C and I-J). In contrast, mutants lacking the PAM2 domain remained cytoplasmic after stress in both S2 (Figure 4D) and U2OS cells (Figure 4K). Notably, point mutations in the Ataxin-2-PAM2 domain that specifically disrupt PAM2:PABP interaction similarly prevent localization to stress granules (Figure 4E-G and L-N). Thus, interactions between Ataxin-2’s PAM2 domain and PABP appear important for the presence of Ataxin-2 in native mRNP granules, whose assembly is driven by the distinct (IDR) region of the protein (Figure 2C). The ability of otherwise full-length Ataxin-2 lacking PAM2 to form compositionally distinct mRNP assemblies (Figure 2B) suggests that PAM2:PABP binding also serves to limit non-physiological interactions by Ataxin-2 (See Discussion).

### The IDR and PAM2 domains promote and the LSm domain inhibits cytotoxicity in Drosophila neurodegeneration models

Three different Ataxin-2 domain deletions tested showed three distinct effects on mRNP granule assembly in S2 cells. IDR domain deletions prevent Ataxin-2 granule formation. LSm-domain deletion enhances the formation of Ataxin-2 granules. PAM2 domain deletions result in the formation of unusual mRNP assemblies (Bakthavachalu *et al*., 2018; Singh *et al*., 2021) (Figure 2B/C). Prior observations showing that Atx2 IDR deletions suppress cytotoxicity in *Drosophila* models for neurodegeneration indicate that mRNP granules support events that lead to degenerative disease (Bakthavachalu *et al*., 2018; Becker *et al*., 2017; Huelsmeier *et al*., 2021; Scoles *et al*., 2017). If true, the expression of Atx2ΔLSm, which enhances granule assembly, would promote or potentially accelerate the degeneration, while the expression of Atx2ΔcIDR would not. The expression of Atx2ΔPAM2 would be expected to support mRNP assemblies of different compositions from the ones containing wild-type or ΔLSm forms of Atx2. The effects on degeneration for this condition would be hard to predict.

To examine how the different Atx-2 domain deletions affect nervous system integrity and function over time, we combined a Gal4-responsive *UAS-Atx2* transgene with *elav-Gal4* and *TubGal80^ts^*. This allows us to use a temperature shift from 18°C to 30°C to induce *UAS-Atx2* transgene expression, specifically in the brains of adult flies (Figure 5A). We then analysed the rate at which flies climbed the walls of a glass cylinder, a surrogate measure of motor ability, one day and 15 days after transgene expression. All genotypes tested showed robust and comparable levels of climbing ability on day 1. Interesting variations were identified on day 15. The 15-day old flies expressing Atx2WT or Atx2ΔLSm showed a strong decline in climbing ability. In contrast, Atx2 ΔcIDR flies showed a minimal decline (Figure 5B). These observations were in line with the effects of these Atx2 types on granule formation. Strikingly, flies expressing the Atx2ΔPAM variant, which formed compositionally distinct granules in S2 cells, showed no significant decline in climbing ability, suggesting that Atx2’s ability to promote progressive decline of neural function depends less on Atx2 granule formation and aggregation, but a bit more on its sequestration of critical translation factors such as PABP (and associated RNAs.(Figure 5B). These observations support and extend prior work showing that heterologous overexpression of full-length, but not PAM2-domain deleted forms of mammalian ATXN2 enhances mammalian TDP-43-induced degeneration of the *Drosophila* compound eye (Kim *et al*., 2014). They are also consistent with work in mice showing that PABPC1 sequestration in inclusions correlates strongly with the progression of neurodegeneration (Damrath *et al*, 2012).

**Figure 5:**
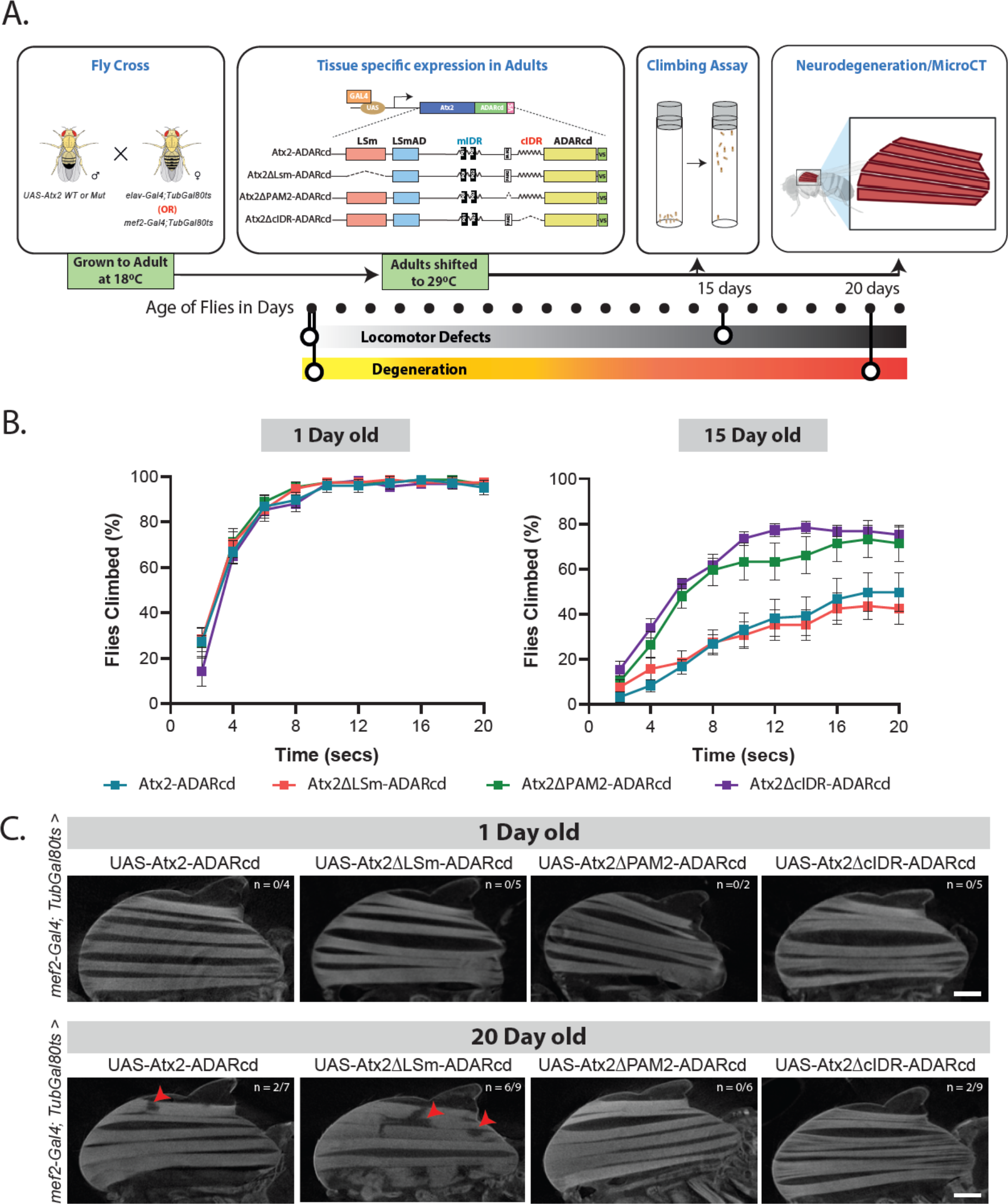
The IDR and PAM2 domains promote and the LSm domain inhibits neurodegeneration in *Drosophila*. (A) A schematic of the experimental design is shown. UAS-transgenes were crossed with *elav-Gal4; tub-Gal80^ts^* or *mef2-Gal4, tub-Gal80^ts^* and kept at 18°C till the adult flies emerged. The flies were shifted to 29 °C for days shown with dots under the experimental design. Fly climbing or indirect flight muscle cytotoxicity was studied. (B) *Drosophila* climbing behavior assay was performed by driving UAS-transgene (Atx2WT, Atx2ΔcIDR, Atx2ΔPAM or Atx2ΔLSm) with *elav-Gal4.* A graph was plotted with number of flies (Y-axis) that crossed the 20ml mark at a given time (X-axis). (C) Cellular toxicity was measured by driving UAS-transgene (Atx2WT, Atx2ΔcIDR, Atx2ΔPAM or Atx2ΔLSm) with *mef2-Gal4.* Fly indirect flight muscles were imaged using micro-CT and the loss of muscle fibers is shown with solid red arrowheads.

The conclusion, that PAM2 mediated interactions were required for progressive cytotoxicity, is further supported by a parallel series of experiments in which we used *mef2-Gal4* in place of *elav-Gal4,* to *target UAS-Atx2* transgene expression to *Drosophila* adult muscles (Figure 5C). Micro-computed tomography (micro-CT) scanning to visualize the integrity of flight muscle fibers in whole-mount preparations (see Methods) revealed degeneration of muscles expressing wild-type Atx2 in 20-day old flies. While there was more severe degeneration in Atx2ΔLSm expressing muscle, muscles similarly expressing Atx2ΔcIDR or Atx2ΔPAM forms showed no morphological defects (Figure 5C and Supplementary Figure 4).

## DISCUSSION

The results described above provide three significant lines of insight. First, they support a detailed model for sequential protein-protein interactions through which Ataxin-2 can modulate different translational states of a single mRNA. Second, they show that the Ataxin-2 polypeptide contains distinct activities that promote or protect against neurodegeneration, pointing to the value of developing therapeutics that target specific Ataxin-2 interactions, beyond those that reduce overall levels of the protein. Third, the work identifies a novel molecular mechanism involving the PAM2 domain and PABP that contributes to the assembly of mRNP granules.

### Molecular mechanisms of Ataxin-2 function

Some RNA-binding proteins can remain associated with mRNAs across multiple stages: RNA processing, transport, translation, or translational control (Formicola *et al*., 2019; Gomes & Shorter, 2019; Hachet & Ephrussi, 2004; Harlen & Churchman, 2017; Lin *et al*, 2015; Maniatis & Reed, 2002). Ataxin-2 may be one such protein. It is a translational activator of the *Drosophila period* mRNA, a repressor of several miRNA reporters, a facilitator of neuronal mRNP-granule and stress-granule formation as well as a broad stabilizer of Ataxin-2 associated mRNAs (Bakthavachalu *et al*., 2018; Inagaki *et al*., 2020; Lim & Allada, 2013; McCann *et al*, 2011; Nonhoff *et al*, 2007; Sudhakaran *et al*, 2014; Yokoshi *et al*, 2014; Zhang *et al*, 2013). While these different functions could represent different modes of engagement with distinct sets of mRNAs, the data are also consistent with another model. Sequential interactions mediated by different protein regions during mRNP modelling allow Ataxin-2 to contribute in multiple ways to translational control to a single mRNA.

Previous work has shown that Atx-2 enhances *period* mRNA translation through a mechanism requiring LSm-domain interactions with a complex of LSM12 and TYF (Twenty Four) proteins associated with the 5’cap of the translating mRNA (Lee *et al*, 2017; Lim & Allada, 2013; Zhang *et al*., 2013). Given considerable supportive evidence for direct binding between the LSm-domain and LSM12, we postulate that LSm-domain-LSM12 interactions occur in translating polysomes (Satterfield & Pallanck, 2006) and contribute to increased efficiency of translation. This proposal is consistent with the observation that the LSm domain opposes the formation of mRNP granules, which usually contain translationally repressed mRNAs (Singh *et al*., 2021).

However, the LSm domain must also contribute to LSM12-independent functions, because while LSm-domain deletions from *Drosophila* Atx2 cause lethality and LSM12 null mutants, while arrhythmic, are viable and fertile (Lee *et al*., 2017). One possibility is that LSm domains additionally contribute, perhaps indirectly, to interactions with the DEAD-box helicase Me31B/DDX6 in a translational repressor complex (Brandmann *et al*, 2018; Lee *et al*., 2017). Thus, we suggest that in the case of actively translating mRNAs, the Atx2 function is driven by LSm-domain association with LSM12 and translational initiators, and that LSM12 disengages from a translational initiation complex as the mRNA transitions into a repressed state driven by Me31B.

While polyA tails and PABP are known to support translation and the Ataxin-2 PAM2 domain is involved in targeting the protein to polysomes (Satterfield and Pallanck, 2006), existing data do not directly address how Ataxin-2 PAM2 motif interactions contribute to translational activation. One possibility, supported by observations on the *period* mRNA is that the PAM2-domain guides Ataxin-2 to the 3’UTR of its target mRNAs (Lim & Allada, 2013). Our observation that PABP co-immunoprecipitates with mini-Ataxin2, show that Atx2-PAM2:PABP interactions occur independently of and prior to mRNP granule formation. Recent findings that this association antagonizes the Ataxin-2 condensation (Boeynaems *et al*., 2021) are consistent with a model in which the Atx2-PAM2 motif interacts with PABP in translating mRNAs to support efficient translation driven by the LSm-LSM12 complex. However, in addition to supporting translation, PABP is also known to associate with translational repressors that could drive either mRNA deadenylation and/or storage (Machida *et al*, 2018; Yoshida *et al*, 2006). Our data support such a dual role for Ataxin-2 associated with PABP in translational repression. First, when Ataxin-2 target mRNAs are not actively translated, then the mRNP through Me31B/DDX6 and PABP may recruit deadenylases to transition into either a translationally dormant or degradative state (Lee *et al*., 2017; Machida *et al*., 2018; Yi *et al*, 2018; Yoshida *et al*., 2006). Second, Atx2 associated mRNA may move into mRNP granules whose formation is facilitated by Atx2 IDR-mediated condensation. We postulate that mRNAs in such assemblies are stored in a form that is protected from degradation. While the above model, shown in Figure 6, is consistent with all our data, we acknowledge that it needs extensive and rigorous testing in the context of the life cycle of a single Ataxin-2 target mRNA.

**Figure 6:**
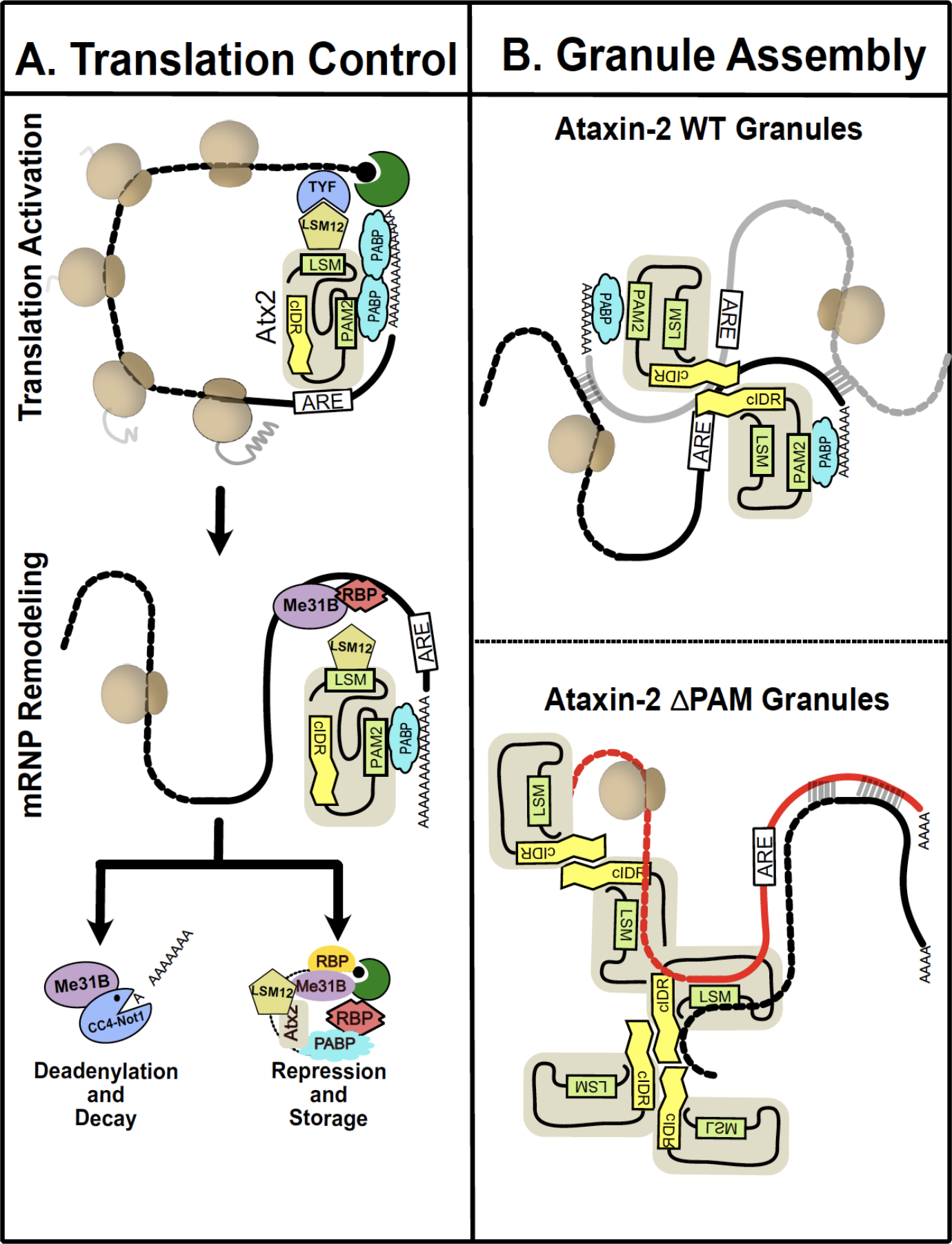
A model for Ataxin-2 RNP dynamics and the role of PAM2 domain in determining its RNP composition and mRNA selection. (A) Ataxin-2 is recruited to mRNAs by RBPs during different stages of the mRNA life cycle. Ataxin-2 activates translation of subsets of mRNA by recruiting LSM12, TYF and other translation activation complexes. Under specific conditions, mRNP remodelling exposes Ataxin-2 cIDR that mediates multivalent interactions and RNP granule assembly. Ataxin-2 recruits Me31B and CCR4-NOT1 complexes that lead to deadenylating and/or translation repression. It is possible that LSM12/TYF continue to associate with RNA but are probably not part of repressor complexes. RNA deadenylation can lead to degradation or translation repression and storage in RNP granules. (B) Ataxin-2-PAM2 domain determines protein and RNA partners of the RNP granules. PAM2 domain is essential for recruitment of Ataxin-2 to stress granules that also contains other RBPs (eg. Me31B, FMRP, Rox8, Rin and Caprin). Ataxin-2-cIDR along with RNA-RNA interaction stabilise the stress induces RNP condensation. In the absence of the PAM2 domain, Ataxin-2 fails to recruit specific target mRNA and proteins. Remodelling of Ataxin-2 exposes the cIDR to induce phase separation and aberrant RNP condensation. The Ataxin-2ΔPAM2 granules are non-toxic and lack several known stress granule proteins (eg.FMRP, Caprin and PABP).

### Implications for Ataxin-2 as a therapeutic target

Antisense Oligonucleotide (ASO) based therapeutic strategies that lower levels of Atxn-2 are being developed for the treatment of ALS and spinocerebellar ataxia type 2 (SCA2) (Becker *et al*., 2017; Scoles *et al*., 2017). Our experiments provide a much finer grained analysis of activities of Ataxin-2, suggesting that the function of the LSm domain should be spared, and that IDR mediated assembly mechanisms and perhaps PAM2:PABP interactions should be most usefully targeted by therapeutics.

Our previous work showed that *Atx2* mutants lacking the cIDR required for Ataxin-2 granule formation in Drosophila neurons and S2 cells, were resistant to neurodegeneration as assessed in *Drosophila* disease models (Bakthavachalu *et al*., 2018; Huelsmeier *et al*., 2021). We further showed that the LSm-domain antagonizes Ataxin-2 granule formation (Singh et al, 2021). Here we advance the latter observation by demonstrating that Ataxin-2 forms lacking the LSm domain may more effectively cause cytotoxicity than the wild-type or IDR-deficient forms (Figure 5C). These observations independently confirm our original conclusions and provide further support for a model in which the efficiency of mRNP assembly correlates with the speed and severity of neurodegenerative processes in *Drosophila*.

The importance of the PAM2 domain in promoting degeneration has been previously observed by experiments showing that heterologous expression of a pathogenic form of human Ataxin-2 lacking its PAM2 domain, but not the full-length form, suppresses cytotoxicity in *Drosophila* expressing human TDP-43 (Kim *et al*., 2014). Our observations that expression of Atx2ΔPAM2 is far less toxic than expression of wild-type Atx2 is consistent with this. In addition, by showing that Atx2ΔPAM2 forms compositionally different Ataxin-2 granules, they highlight the importance of specific granule components, and not granules *per se,* in neurodegenerative pathologies. Thus, while liquid-liquid transitions mediated by disordered domains could be a shared requirement for the formation of multiple types of mRNP granules, we speculate that each granule type, with distinctive composition, could preferentially support one or other type of proteinopathy (De Graeve & Besse, 2018; Vogler *et al*, 2018).

### Structured interactions may determine mRNP granule composition

Many lines of evidence argue that specific molecular interactions, e.g. mediated by structured domains of the P-body component Edc3 or the stress-granule components G3BP and Caprin, contribute to the mRNP granule formation (Decker *et al*., 2007; Kedersha *et al*, 2016). In engineered systems, the condensation of RNA-binding proteins and mRNAs into granules has been clearly shown to depend on both traditional protein-protein interactions and on more promiscuous interactions between intrinsically disordered regions (Protter *et al*, 2018). Our work now identifies the interactions between Ataxin-2’s PAM2 motif and PABP as a critical contributor to the assembly of Ataxin-2 containing mRNP granules. This suggests a mechanism by which the interaction helps select mRNA and protein components of mRNP granules.

We suggest that Ataxin-2, guided by PAM2:PABP interactions and LSm domain interactions, recruits target mRNAs and associated proteins into translating mRNPs (Satterfield & Pallanck, 2006). Under conditions where the translation is arrested, LSm-domain interactions are altered (Lee *et al*., 2017), and transcripts are released from stalled ribosomes. Base-pairing interactions between exposed mRNA side chains, as well as interactions between Ataxin-2’s now accessible intrinsically disordered regions, contribute to the assembly of these mRNPs into granules. This logical sequence of events is consistent with: (a) TRIBE data showing a reduced number of edits of native Ataxin-2 target mRNAs by Atx2ΔPAM2-ADARcd; (b) the inability of ΔPAM2-miniAtx2 constructs to associate with stress granules; and (c) the aberrant protein composition of granules induced by Atx2ΔPAM2 in S2 cells. The additional observation that Atx2ΔPAM2-ADARcd expression results in a large number of non-native mRNA edits, indicates that the PAM2:PABP interaction not only selects correct target mRNAs but also prevents Ataxin-2 engagement with incorrect mRNA target regions.

Our conclusion that Ataxin2-PAM2:PABP interactions are involved in the selection of mRNA components of RNP granules is superficially inconsistent with the observation that RNA components of native stress granules can be predicted with remarkable accuracy on the basis of mRNA size. This argues for a primary role for RNA-RNA interactions in the stress granule assembly (Jain & Vale, 2017; Matheny *et al*., 2021; Van Treeck & Parker, 2018). However, we note that experiments presented here do not address mechanisms by which mRNAs are selected into stress granules. Instead, the TRIBE data address how Atx2-target mRNAs are selected into neuronal mRNP granules that exist in non-stressed cells *in vivo*, and microscopic studies analyse protein components of mRNP granules formed following Atx2 expression in S2 cells. Our experiments and observations therefore point to fundamental differences in mechanisms by which the assembly of neuronal granules, or granule types found in unstressed cells, may differ from those involved in stress-granule assembly. The regulation and composition of the former class could well rely extensively on specific protein-protein and protein-mRNA interactions, which may be revealed by future analyses of mechanisms by which such mRNP assemblies are formed *in vivo*.

## Materials and methods

### Key resources table

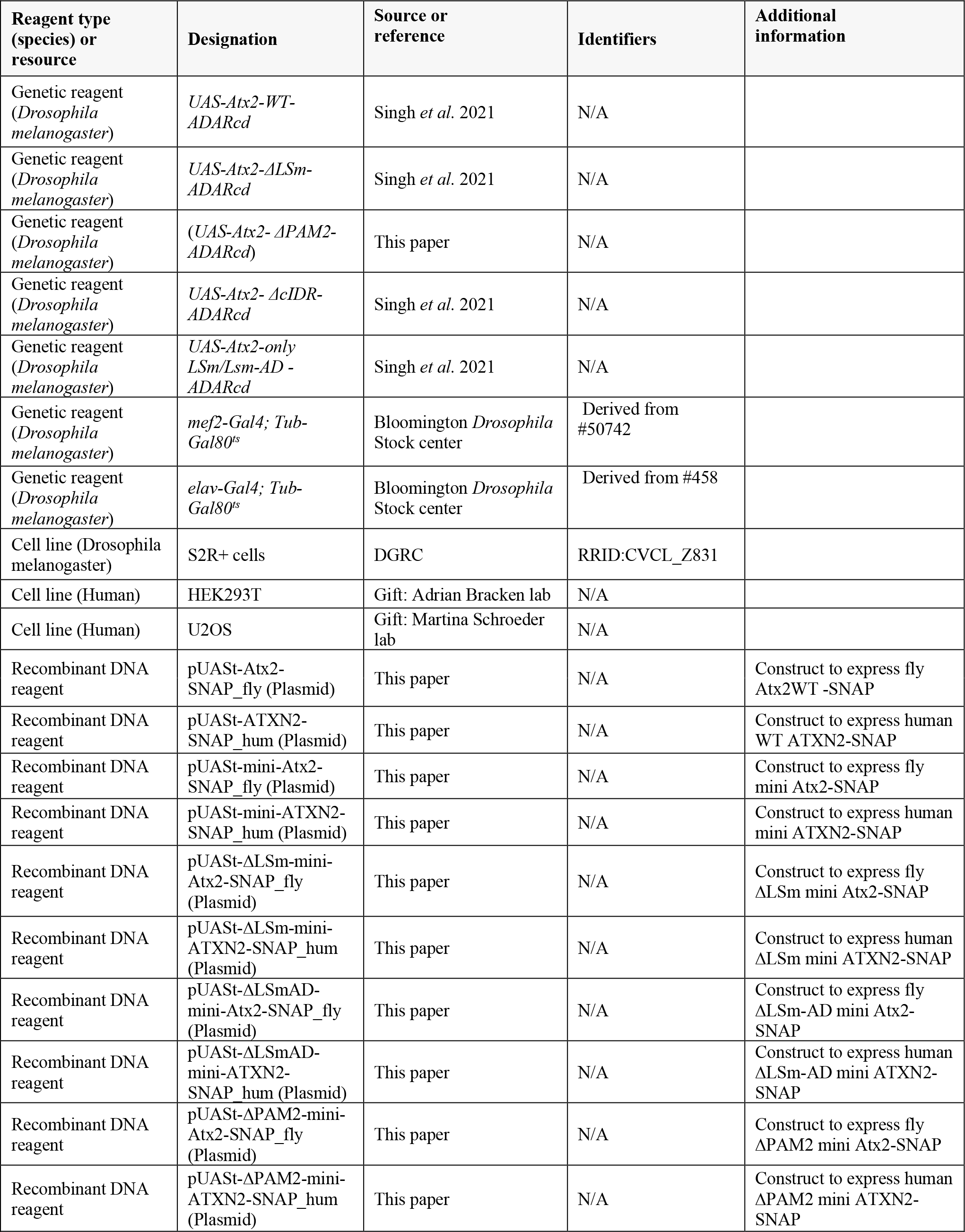
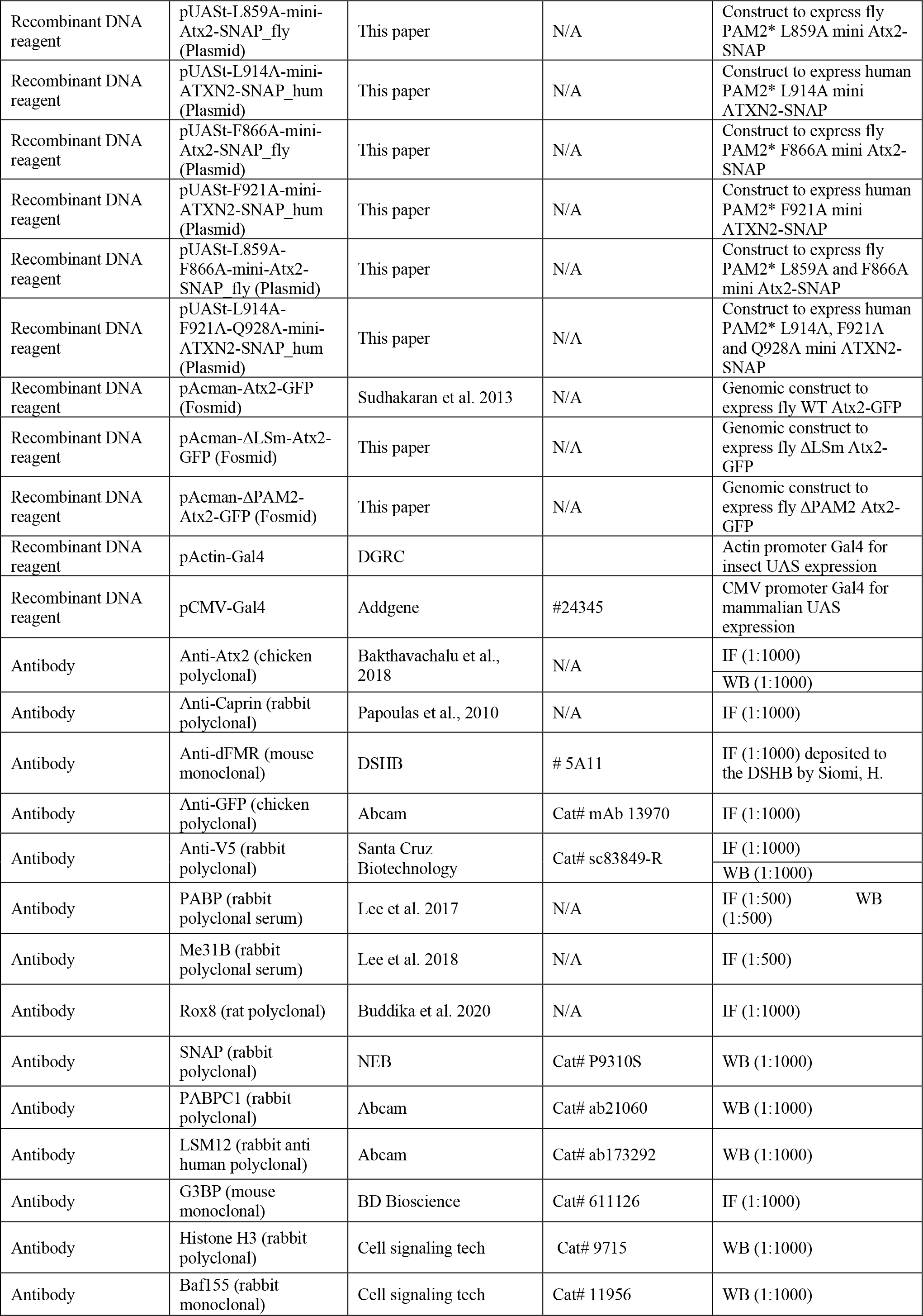
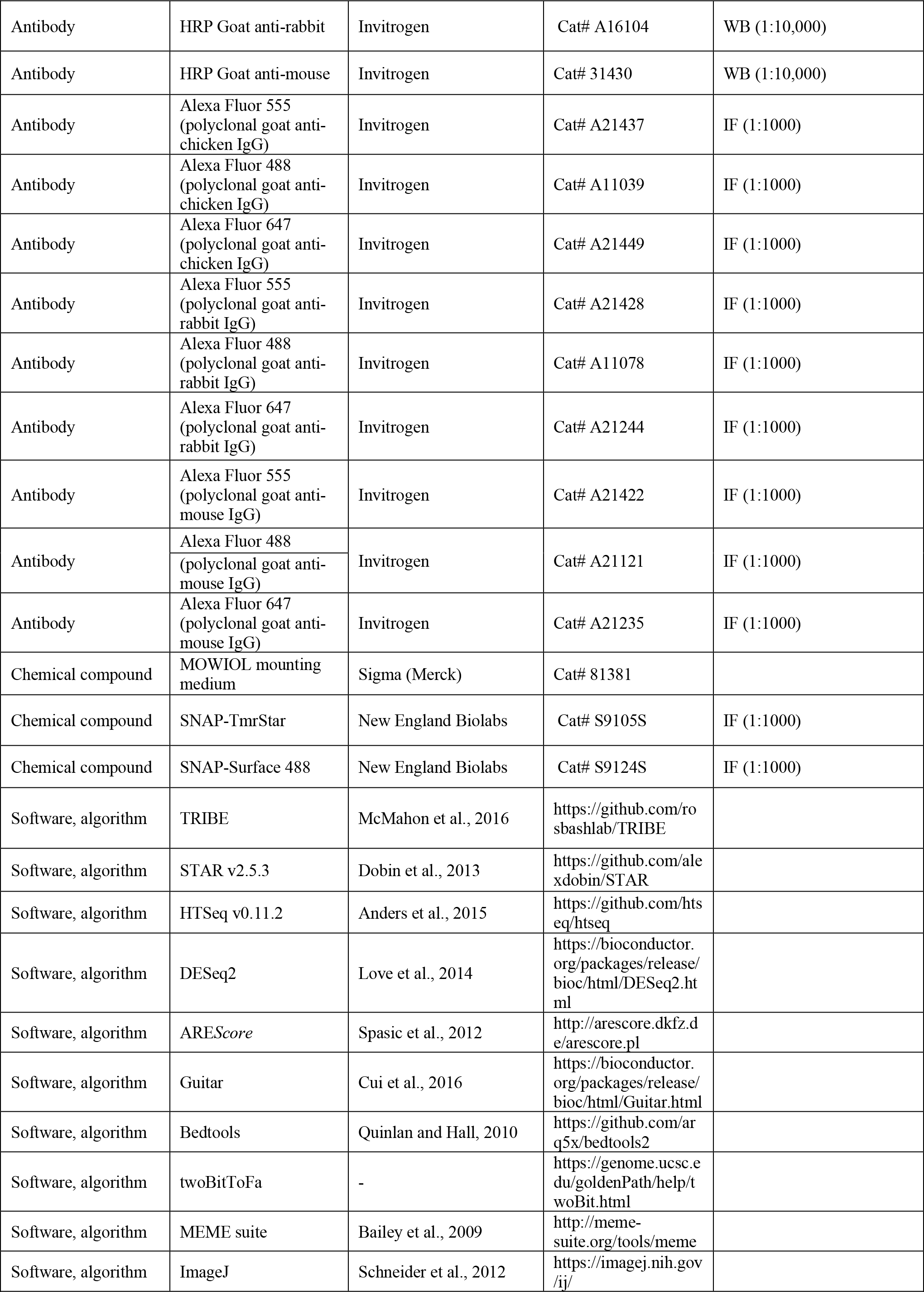
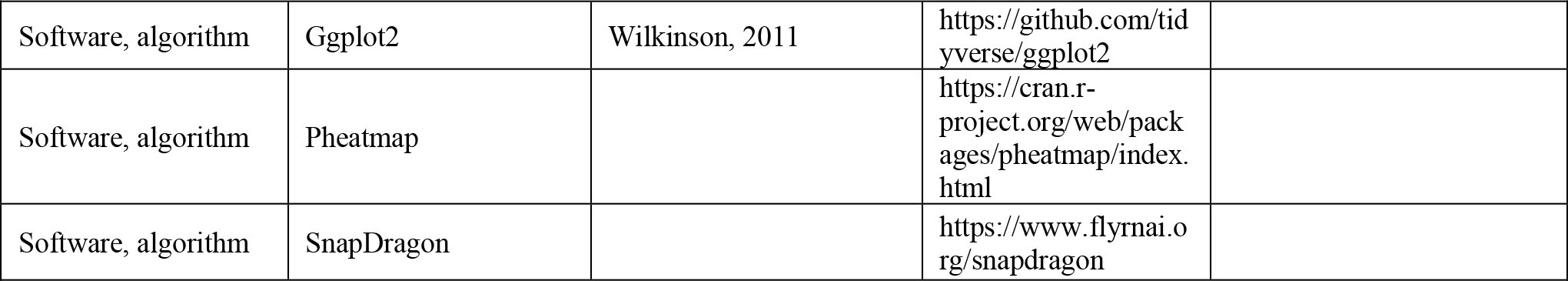

### Cell culture, transfection and stress induction

*Drosophila* S2R+ cells were obtained from the DGRC, Indiana University, and were grown in Gibco Schneider’s S2 media with 10% FBS and 1% penicillin and streptomycin, at 25°C. Transfections were performed using either FugeneHD (Active Motif) or TransIT-X2 (Mirus) reagents at 2:1 ratio μl reagent to μg plasmid DNA for 24-72 h depending on downstream use. HEK293T cells from Adrian Bracken, Trinity College Dublin, were grown in Gibco Dulbecco’s Modified Eagle Media with 10% FBS, 2 mM l-glutamine addition, 1% penicillin and streptomycin, at 37°C and 5% CO2. U2OS cells from Martina Schroeder, Maynooth University, were grown at the same conditions as HEK293T. Mammalian cell transfections were carried out with 1 mM PEI (Polysciences) solution at 2:1 ratio μl reagent to μg plasmid DNA for 24-72H depending on downstream use. For confocal imaging applications cells were grown in 24-well plates on glass cover-slips for 24 h before transfection for up to 48 h. For Western blotting and IP, cells were grown in 75 cm^2^ flasks until >80% confluent before transfection for up to 72H before harvesting. Oxidative stress was induced in *Drosophila* S2R+ cells with addition of sodium arsenite solution to a final concentration of 50 μM in media for 3 h. In mammalian cells, oxidative stress was induced in the same way except for only 1 h.

### Western blotting and protein immunoprecipitation

Total protein extracts were prepared from S2 and HEK293 cells as described earlier (Sudhakaran et al., 2014). Up to 10 µg total protein was loaded per well for detecting Atx2-SNAP constructs, partner proteins and loading controls on 8-12% SDS-PAGE gels and transferred to nitrocellulose membranes. The blots were probed in 5% skim milk in PBS using rabbit anti-SNAP (1:1000), rabbit anti-PABP (1:1000), rabbit anti-LSM12 (1:1000) antibodies, and mouse anti-histone H3 (1:5000) and mouse anti-BAF155 (1:2000) loading control antibodies. Corresponding HRP-conjugated secondary antibodies were used at 1:10,000 dilution and developed using Pierce ECL western blotting substrate (ThermoFisher) as per the manufacturer’s instructions.

For Atx2-SNAP construct immunoprecipitation, transfected cell lysates were normalised to the same volume and concentration, 10% of the volume was saved and diluted as an input control, and Chromotek anti-SNAP-tag conjugated agarose beads and IP kits were used according to the manufacturer’s specifications. Pulled-down proteins together with corresponding sample input controls were blotted as described above.

### Immunohistochemistry and imaging of cultured cells

Transfected cells on coverslips were fixed with 4% paraformaldehyde in PBS solution for 15 min, followed by three 5 min washes in PBS. Permeabilization was performed on all cells with 0.5% TritonX100 in PBS solution for 3 min, before three more 5 min washes in PBS. Cells were blocked with 3% BSA in PBS for 1 h at room temperature before staining with primary antibodies at appropriate dilutions in 3% BSA overnight at 4°C. Corresponding fluorescent secondary antibodies in 3% BSA were used to stain the sample cells for 1 h at room temperature after primaries were washed off. Where SNAP-tagged proteins were being visualized, SNAP-ligand TMR-Star (NEB) or SNAP-surface-Alexa488 (NEB) were added at the secondary antibody staining stage. Following staining and washing, cells were mounted upside-down on microscopy slides in MOWIOL, allowed to cure for >12 h at 4 °C, and imaged on a Zeiss LSM880 Airyscan/AiryscanFast confocal microscope with a 20x air objective.

### Bioimage analysis

Where relevant, Airyscan images were processed with Zen Black software (Zeiss) with recommended settings. Confocal microscopy images were analysed using macros within ImageJ/FIJI and Excel. Quantification of co-localisation was performed by comparing stress granule marker staining intensity profiles across a randomised selection of Atx2 granules within transfected cells, with the intensity profile of the Atx2 staining. Any signal 10% or higher than background (adjusted for fluorophore bleed through) was deemed evidence of co-localisation within that particular granule. For quantifying the exclusion of mini-Atx2-SNAP constructs from stress induced granules the Caprin or G3BP1 staining was used as independent identifier of stress granules and Atx2 profiles were compared to them. 48-120 granules were quantified in each co-staining (Figure 2), and 28-70 granules were quantified for each construct transfection (Figure 4).

### Crystal structure threading

Threading of the *Drosophila* PAM2 peptide bound to the MLLE domain of PABPC1 was performed using the Swiss-PdbViewer software, based on the human crystal structure of the complex obtained from PDB, identified as 3KTR (Kozlov et al., 2010)

### Experimental fly crosses

*Drosophila* stocks were maintained at 25°C in corn meal agar. Strains homozygous for UAS-transgenes were crossed with *elav-Gal4* and *tub-Gal80ts* at 18 °C till the adult fly emerged. The flies were shifted to 29 °C for 5 days before processing for RNA extraction for TRIBE experiments. The climbing behaviour experiments were performed on flies kept at 29 °C for either 1 or 15 days. For microCT experiments, the UAS-transgenes were crossed with *mef2-Gal4* and *tub-Gal80ts* at 18°C and the adult flies were transferred to 29 °C for 1 day or 20 days.

### RNA extraction from brain and NGS

Around 10-12 adult brains were dissected in RNA Later for total RNA isolation. RNA was isolated using TRIzol reagent (Invitrogen) as per the manufacturer’s protocol. Poly(A)-enriched mRNA was used to prepare Illumina libraries using the NEBNext Ultra II Directional RNA Library Prep kit (E7765L). Atx2-ΔPAM2-ADARcd samples were sequenced with Illumina HiSeq PE Rapid Cluster Kit v2 (PE-402-4002) to generate 2 × 100 paired-end strand-specific data using the Illumina HiSeq 2500 sequencing platform.

### TRIBE data analysis

The sequencing reads obtained had a mean quality score (Q-Score) >= 37. Analysis of the TRIBE data was performed as described previously (McMahon et al., 2016, Singh et al., 2021). Briefly, the reference genome and gtf file of *Drosophila melanogaster*, version dm6, were downloaded from the UCSC genome browser. Raw sequencing reads were mapped using TopHat2 (Trapnell *et al*, 2009) with the parameters ‘--library-type fr-firststrand -m 1 N 3 -- read-edit-dist 3 p 5 g 2 -I 50000 --microexon-search --no-coverage-search -G dm6_genes.gtf’. Only uniquely mapped reads are considered for editing analysis. A table of raw and mapped reads is included in Supplementary table 1. A threshold file was created by ensuring only edits with coverage of at least 20 reads and 15% edits were retained. All the TRIBE experiments were performed in duplicates, and only the edits identified in both replicates above the edit threshold are reported.

### Climbing Assay

Appropriately aged adult *Drosophila* was transferred to a 50 ml graduated glass measuring cylinder for the climbing assay and sealed with a cotton plug. A digital video camera was positioned to record the vials. The assay was initiated by tapping the cylinder against a foam pad to collect the flies to the bottom of the cylinder and the flies were allowed to climb the cylinder with video being recorded for ∼30 s. The number of flies that crossed the 20 ml mark (∼5.5cm) was counted over time and the data was plotted against the time using GraphPad prism. Average of 3 trials were used for each biological replicate. 7-10 biological replicates were used for each genotype.

### Sample preparation and scanning for microCT

*Drosophila* indirect flight muscle microCT was carried out as described in Chaturvedi et. al, 2019. Briefly, animals were anesthetized on ice and fixed in PBS containing 4% paraformaldehyde (PFA). Thoraces were dissected and stained using 1% elemental iodine (1.93900.0121, Emparta, Merck) with 2% potassium iodide (no. 15 724, Qualigens) dissolved in PBS. The stained samples were washed in PBS and embedded in petroleum jelly. MicroCT scanning was carried out at 40 kV, 250 µA, on Bruker Skyscan-1272.

### Data availability

The RNA sequencing data have been deposited to GEO under the accession code GSE196739.

### Contact for reagent and resource sharing

Further information and requests for resources and reagents should be directed to and will be fulfilled by the lead contacts Mani Ramaswami and Baskar Bakthavachalu.

## ACKNOWLEDGEMENTS

We thank members of the Ramaswami, Vijay Raghavan and Bakthavachalu labs and Roy Parker for useful discussions and/or comments on the manuscript. We thank Marlena Mucha, Adrian Bracken and Amir Khan for advice and help with biochemical experiments, Michael Rosbash for *Drosophila* TRIBE plasmids, and colleagues acknowledged in the Key Resources table for reagents and advice. The fly facility at Bangalore Life Science Cluster (BLiSC) provided support with fly stock supply as well as generation of transgenic; CIFF at BLiSC provided essential confocal microscopy support; and Awadhesh Pandit and next-generation genomics facility at BLiSC provided NGS service. Daniel Fortunati thanks Kenneth Mok for his mentorship. We acknowledge Drosophila Genomics Resource Centre (supported by NIH grant 2P40OC010949) for *Drosophila* S2 cells.

## AUTHOR CONTRIBUTIONS

Conceptualization, A.P., D.F, A.S., J.Huelsmeier, K.V.R., M.R., and B.B.; Methodology, A.P., D.F, A.S., J.Huelsmeier, A.R.K., S.S.P., J.Hillebrand, K.A., D.J., G.B., J.L., C.L., G.A., K.H.M., K.V.R., M.R., and B.B.; Investigation, A.S., J.Huelsmeier, A.R.K., S.S.P., J.Hillebrand, A.P., K.A., K.V.R., M.R., and B.B.; Writing–Original Draft, A.P., D.F, A.S., J.Huelsmeier, K.V.R., M.R., and B.B.; Writing–Review & Editing, A.P., D.F, A.S., J.Huelsmeier, A.R.K., S.S.P., J.Hillebrand, K.A., D.J., G.B., J.L., C.L., G.A., K.H.M., K.V.R., M.R., and B.B.; Funding Acquisition, K.V.R., M.R., and B.B.; Resources, Fly community.

## SUPPLEMENTARY FIGURES

**Supplementary Figure 1:**
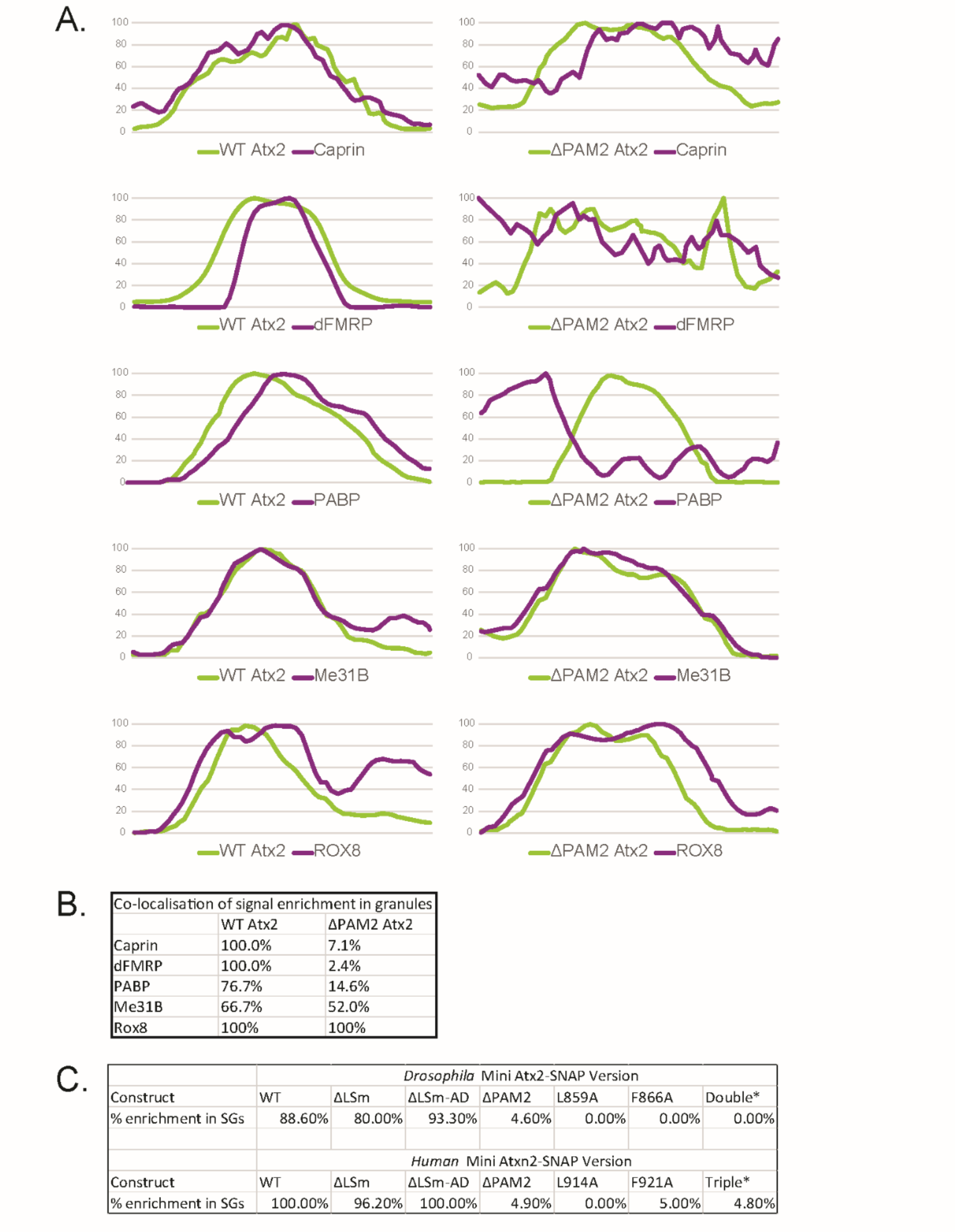
Co-localisation quantification for figure 2, figure 4. (A) Normalised profile plots of Atx2-GFP granules in S2 cells as shown in Figure 2. Within representative granules of wild type Atx2-GFP (green line), SG components Caprin, dFMRP, PABP, Me31B, and Rox8 show largely overlapping enrichment of fluorescence profile along a line bisecting a granule after immunohistochemistry and imaging (purple line). In Atx2ΔPAM2-GFP granules, this colocalization of fluorescence signals is not seen in the case of Caprin, dFMRP and PABP, suggesting these components are not enriched in these granules above background level. (B) Quantification of co-localization for Figure 2. N = 48-120 images of Atx2-GFP granules were randomly selected for each co-staining and analysed for signal co-enrichment (see methods) in the case of each component assayed. (C) Quantification of Atx2 construct inclusion in SGs for Figure 4. N = 28-70 images of stress granules in arsenite stressed S2 cells (marked by anti-Caprin staining) and U2OS cells (marked by anti-G3BP staining) were randomly selected for each Atx2 construct assayed and were analysed for Mini Atx2-SNAP allele signal co-enrichment (see methods).

**Supplementary Figure 2:**
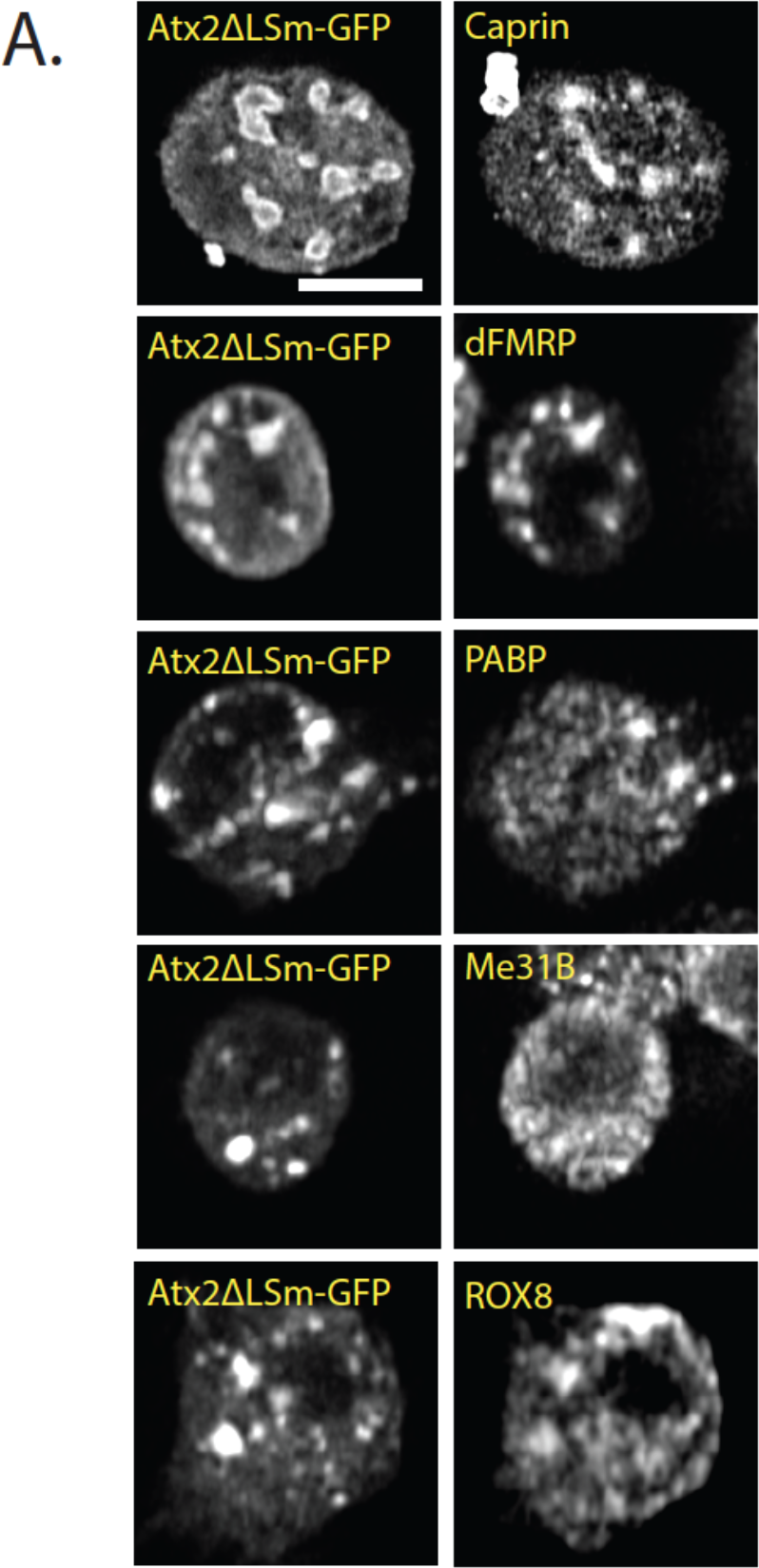
Atx2ΔLSm granules in S2 cells do not show significantly altered protein contents compared to wild-type Atx2. Caprin, dFMRP, PABP, Me31B, and Rox8 co-localize with overexpressed Atx2ΔLSm GFP, suggesting that the granules formed contain a similar set of components as Atx2 granules. It should be noted that Atx2 granules do not sequester the majority of the endogenous components stained for, leading to a high, diffuse background staining.

**Supplementary Figure 3:**
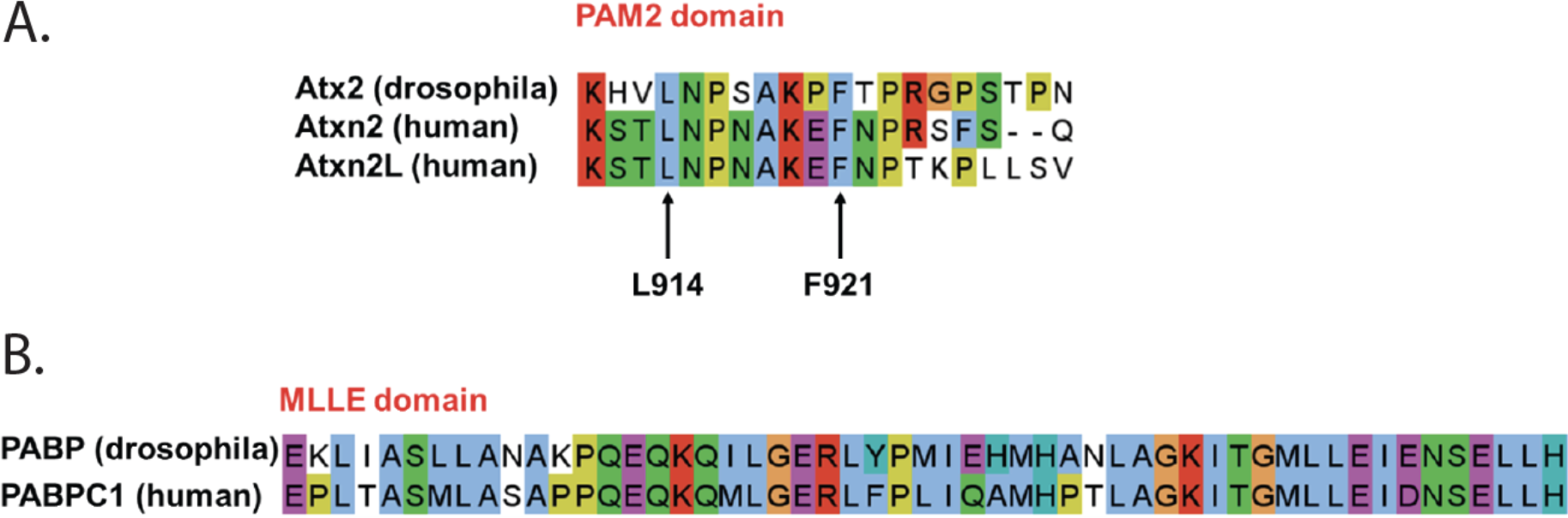
The ATXN2 PAM2 and the PABPC1 MLLE domain are highly conserved from fly to human. (A) The ATXN2 PAM2 domain exhibits high sequence similarity where the key MLLE domain hydrophobic binding residues leucine 914 and phenylalanine 921 (human ATXN2 numbering) are conserved from *Drosophila* to humans. (B) Its binding partner, the PABPC1 MLLE domain, is also highly conserved from *Drosophila* to human. Sequence IDs: Q8SWR8 (Atx2_DROME), Q99700 (ATXN2_HUMAN), Q8WWM7 (ATX2L_HUMAN), P21187 (PABP_DROME), P11940 (PABP1_HUMAN).

**Supplementary Figure 4:**
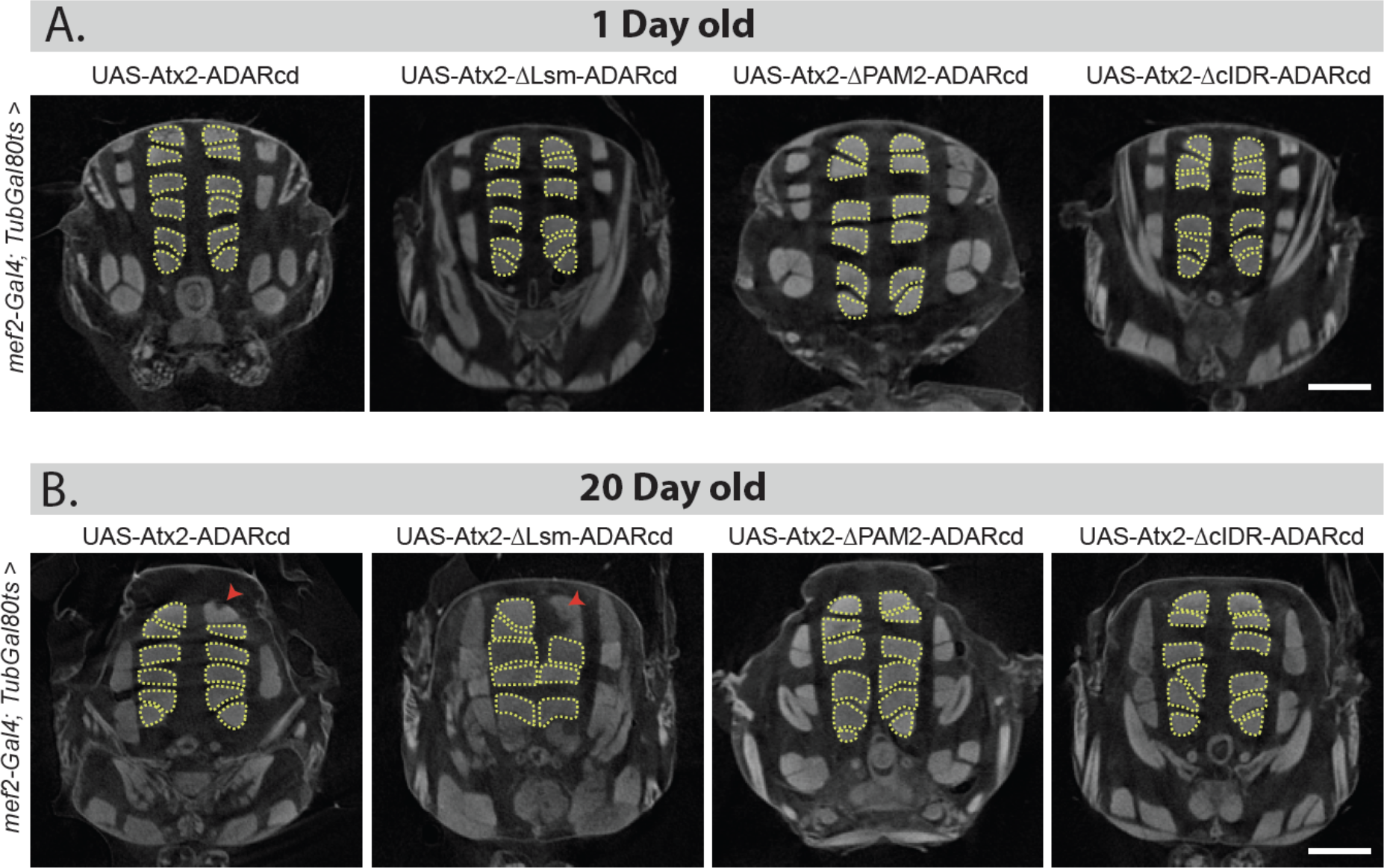
Transverse view of *Drosophila* indirect flight muscle imaged using micro-CT show cellular toxicity. (A) Driving UAS-transgene (Atx2WT, Atx2ΔcIDR, Atx2ΔPAM or Atx2ΔLSm) with *mef2-Gal4* show normal muscles on day 1. (B) Expression of wild-type and Atx2ΔLSm transgene for 20 days show loss of muscle fibers, indicated with solid red arrowheads. Expression of Atx2ΔPAM2 and Atx2ΔcIDR for 20 days show no visible phenotype.

**Supplementary Table 1:**
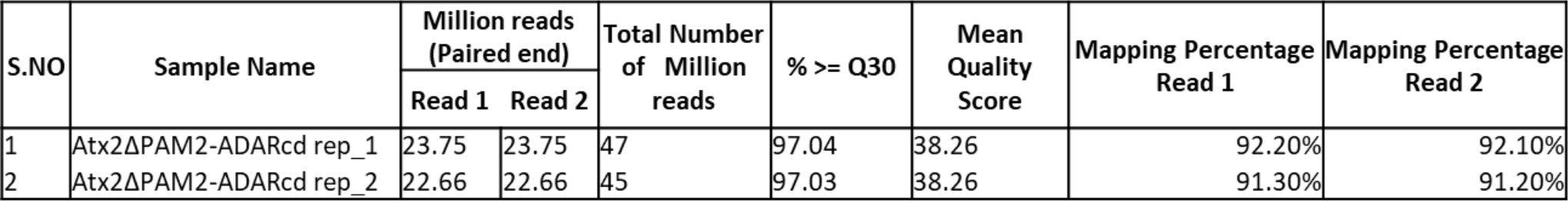

| S.NO | Sample Name | Million reads (Paired end) |  | Total Number of Million reads | % >= Q30 | Mean Quality Score | Mapping Percentage Read 1 | Mapping Percentage Read 2 |
| --- | --- | --- | --- | --- | --- | --- | --- | --- |
|  |  | Read 1 | Read 2 |  |  |  |  |  |
| 1 | Atx2ΔPAM2-ADARcd rep_1 | 23.75 | 23.75 | 47 | 97.04 | 38.26 | 92.20% | 92.10% |
| 2 | Atx2ΔPAM2-ADARcd rep_2 | 22.66 | 22.66 | 45 | 97.03 | 38.26 | 91.30% | 91.20% |

**Supplementary Table 2:** The targets common between Atx2 wild-type and del-PAM2 are shown in bold text.

| Chr | Start | End | Genes | Replicate 1 edit percentage | Replicate 2 edit percentage | Average edit percentage | Chr coordinate |
| --- | --- | --- | --- | --- | --- | --- | --- |
| chr2L | **1009080** | **1009081** | **IA-2** | **17.1** | **15.5** | **16.3** | **chr2L_1009081_IA-2** |
| chr2L | **1009440** | **1009441** | **IA-2** | **31.7** | **38.8** | **35.25** | **chr2L_1009441_IA-2** |
| chr2L | **1011482** | **1011483** | **IA-2** | **31.7** | **34.7** | **33.2** | **chr2L_1011483_IA-2** |
| chr2L | 13505104 | 13505105 | B4 | 22.4 | 17.48 | 19.94 | chr2L_13505105_B4 |
| chr2L | **17188307** | **17188308** | **beat-IIIc** | **18** | **21.7** | **19.85** | **chr2L_17188308_beat-IIIc** |
| chr2L | 19746161 | 19746162 | CG10631 | 25.65 | 19.4 | 22.52 | chr2L_19746162_CG10631 |
| chr2L | **20056018** | **20056019** | **sNPF** | **16** | **24.3** | **20.15** | **chr2L_20056019_sNPF** |
| chr2L | 7891375 | 7891376 | Snoo | 18.5 | 16.7 | 17.6 | chr2L_7891376_Snoo |
| chr2L | 8113160 | 8113161 | Bsg | 17.4 | 19.05 | 18.23 | chr2L_8113161_Bsg |
| chr2L | **8113894** | **8113895** | **Bsg** | **16.82** | **28.7** | **22.76** | **chr2L_8113895_Bsg** |
| chr2L | 8116005 | 8116006 | Bsg | 18.2 | 27.4 | 22.8 | chr2L_8116006_Bsg |
| chr2L | 9256901 | 9256902 | Ggamma30A | 19.5 | 32.6 | 26.05 | chr2L_9256902_Ggamma30A |
| chr2L | **9292530** | **9292531** | **Ggamma30A** | **42** | **54.4** | **48.2** | **chr2L_9292531_Ggamma30A** |
| chr2L | **9292829** | **9292830** | **Ggamma30A** | **42.6** | **53.55** | **48.08** | **chr2L_9292830_Ggamma30A** |
| chr2L | 9295031 | 9295032 | Ggamma30A | 23.5 | 26 | 24.75 | chr2L_9295032_Ggamma30A |
| chr2R | 12085463 | 12085464 | jeb | 16 | 28.4 | 22.2 | chr2R_12085464_jeb |
| chr2R | **13513577** | **13513578** | **Vmat** | **64.2** | **63.9** | **64.05** | **chr2R_13513578_Vmat** |
| chr2R | 13513654 | 13513655 | Vmat | 27.7 | 33.45 | 30.58 | chr2R_13513655_Vmat |
| chr2R | **13513732** | **13513733** | **Vmat** | **24** | **29.75** | **26.88** | **chr2R_13513733_Vmat** |
| chr2R | **13513762** | **13513763** | **Vmat** | **44.55** | **47.3** | **45.92** | **chr2R_13513763_Vmat** |
| chr2R | **13514350** | **13514351** | **Vmat** | **37.7** | **38.5** | **38.1** | **chr2R_13514351_Vmat** |
| chr2R | 13519080 | 13519081 | Vmat | 18.1 | 20.8 | 19.45 | chr2R_13519081_Vmat |
| chr2R | **23690701** | **23690702** | **Pal2** | **17.3** | **17.2** | **17.25** | **chr2R_23690702_Pal2** |
| chr2R | 24213943 | 24213944 | CG30419 | 27.8 | 22.98 | 25.39 | chr2R_24213944_CG30419 |
| chr2R | 24214234 | 24214235 | CG30419 | 24.2 | 26.1 | 25.15 | chr2R_24214235_CG30419 |
| chr2R | 24214648 | 24214649 | CG30419 | 16.4 | 15.9 | 16.15 | chr2R_24214649_CG30419 |
| chr2R | 24215193 | 24215194 | CG30419 | 43.5 | 29.7 | 36.6 | chr2R_24215194_CG30419 |
| chr2R | 24229341 | 24229342 | CG30419 | 20.8 | 16.3 | 18.55 | chr2R_24229342_CG30419 |
| chr2R | 6914804 | 6914805 | CG30158 | 16.4 | 16 | 16.2 | chr2R_6914805_CG30158 |
| chr2R | **6920815** | **6920816** | **CG30158** | **33.92** | **40.48** | **37.2** | **chr2R_6920816_CG30158** |
| chr2R | **6921435** | **6921436** | **CG30158** | **17.5** | **41.9** | **29.7** | **chr2R_6921436_CG30158** |
| chr2R | **6921976** | **6921977** | **CG30158** | **36** | **40.2** | **38.1** | **chr2R_6921977_CG30158** |
| chr2R | **7718070** | **7718071** | **CG18812** | **15.9** | **19.7** | **17.8** | **chr2R_7718071_CG18812** |
| chr2R | 9473772 | 9473773 | Camta | 21.75 | 30.18 | 25.96 | chr2R_9473773_Camta |
| chr2R | 9479421 | 9479422 | Camta | 23.1 | 26 | 24.55 | chr2R_9479422_Camta |
| **chr2R** | **9480019** | **9480020** | **Camta** | **32** | **41** | **36.5** | **chr2R_9480020_Camta** |
| **chr2R** | **9910328** | **9910329** | **FMRFa** | **42.5** | **25** | **33.75** | **chr2R_9910329_FMRFa** |
| **chr3L** | **11498209** | **11498210** | **chrb** | **15.82** | **20.5** | **18.16** | **chr3L_11498210_chrb** |
| chr3L | 12267895 | 12267896 | CG32100 | 20 | 16.1 | 18.05 | chr3L_12267896_CG32100 |
| chr3L | 1504295 | 1504296 | Psa | 18.6 | 15.9 | 17.25 | chr3L_1504296_Psa |
| chr3L | 1521650 | 1521651 | Psa | 18.5 | 34.6 | 26.55 | chr3L_1521651_Psa |
| chr3L | 1543675 | 1543676 | CG7852 | 15.5 | 22 | 18.75 | chr3L_1543676_CG7852 |
| chr3L | 17062355 | 17062356 | Rbp6 | 15.2 | 18.5 | 16.85 | chr3L_17062356_Rbp6 |
| chr3L | 17147382 | 17147383 | Rbp6 | 18.23 | 26.62 | 22.42 | chr3L_17147383_Rbp6 |
| **chr3L** | **17345219** | **17345220** | **Mip** | **29.9** | **27.2** | **28.55** | **chr3L_17345220_Mip** |
| **chr3L** | **17345290** | **17345291** | **Mip** | **17.9** | **16.9** | **17.4** | **chr3L_17345291_Mip** |
| **chr3L** | **19066983** | **19066984** | **Mkp3** | **18.9** | **15.7** | **17.3** | **chr3L_19066984_Mkp3** |
| chr3L | 21494821 | 21494822 | Hr78 | 27.1 | 18.4 | 22.75 | chr3L_21494822_Hr78 |
| chr3L | 21831417 | 21831418 | CG7148 | 15.8 | 28.6 | 22.2 | chr3L_21831418_CG7148 |
| chr3L | 21930851 | 21930852 | mub | 15 | 17.2 | 16.1 | chr3L_21930852_mub |
| chr3L | 21931110 | 21931111 | mub | 15.1 | 20.1 | 17.6 | chr3L_21931111_mub |
| chr3L | 22061206 | 22061207 | Oct-TyrR | 15.4 | 16.7 | 16.05 | chr3L_22061207_Oct-TyrR |
| chr3L | 22877661 | 22877662 | Chro | 35.3 | 45.7 | 40.5 | chr3L_22877662_Chro |
| chr3L | 23148124 | 23148125 | CG32350 | 27.4 | 35.3 | 31.35 | chr3L_23148125_CG32350 |
| chr3L | 23747549 | 23747550 | CG17698 | 40.27 | 28.95 | 34.61 | chr3L_23747550_CG17698 |
| chr3L | 23934990 | 23934991 | CG40470 | 23.5 | 22.05 | 22.77 | chr3L_23934991_CG40470 |
| chr3L | 3910071 | 3910072 | Eip63F-1 | 20 | 24.5 | 22.25 | chr3L_3910072_Eip63F-1 |
| **chr3L** | **3954338** | **3954339** | **CG12605** | **35.2** | **40.8** | **38** | **chr3L_3954339_CG12605** |
| **chr3L** | **3954933** | **3954934** | **CG12605** | **35.42** | **51.1** | **43.26** | **chr3L_3954934_CG12605** |
| chr3L | 3957068 | 3957069 | CG12605 | 21.8 | 26.7 | 24.25 | chr3L_3957069_CG12605 |
| chr3L | 3957671 | 3957672 | CG12605 | 18.9 | 22.2 | 20.55 | chr3L_3957672_CG12605 |
| chr3L | 3961590 | 3961591 | CG12605 | 18.3 | 18.5 | 18.4 | chr3L_3961591_CG12605 |
| chr3L | 3992789 | 3992790 | scrt | 21.08 | 27.28 | 24.18 | chr3L_3992790_scrt |
| chr3L | 4092142 | 4092143 | CG14989 | 15.88 | 18.6 | 17.24 | chr3L_4092143_CG14989 |
| chr3L | 4113123 | 4113124 | Ack | 18.4 | 16.7 | 17.55 | chr3L_4113124_Ack |
| chr3L | 4113297 | 4113298 | Ack | 18.4 | 17.9 | 18.15 | chr3L_4113298_Ack |
| chr3L | 572527 | 572528 | hipk | 29.8 | 31 | 30.4 | chr3L_572528_hipk |
| chr3L | 572530 | 572531 | hipk | 42.7 | 43 | 42.85 | chr3L_572531_hipk |
| **chr3L** | **575712** | **575713** | **hipk** | **36.12** | **52.92** | **44.52** | **chr3L_575713_hipk** |
| **chr3L** | **575753** | **575754** | **hipk** | **31.7** | **49.6** | **40.65** | **chr3L_575754_hipk** |
| chr3L | 576730 | 576731 | hipk | 17.1 | 15.2 | 16.15 | chr3L_576731_hipk |
| chr3L | 577020 | 577021 | hipk | 20 | 16.9 | 18.45 | chr3L_577021_hipk |
| **chr3L** | **577417** | **577418** | **hipk** | **51.42** | **62.12** | **56.77** | **chr3L_577418_hipk** |
| chr3L | 577970 | 577971 | hipk | 21.4 | 25.9 | 23.65 | chr3L_577971_hipk |
| chr3L | 578186 | 578187 | hipk | 18.2 | 24.9 | 21.55 | chr3L_578187_hipk |
| chr3L | 578453 | 578454 | hipk | 25.7 | 32.7 | 29.2 | chr3L_578454_hipk |
| chr3L | 579335 | 579336 | hipk | 18.1 | 17 | 17.55 | chr3L_579336_hipk |
| chr3L | 579634 | 579635 | hipk | 20.5 | 19.8 | 20.15 | chr3L_579635_hipk |
| **chr3L** | **580103** | **580104** | **hipk** | **63.1** | **67.7** | **65.4** | **chr3L_580104_hipk** |
| chr3L | 580500 | 580501 | hipk | 15.6 | 18.1 | 16.85 | chr3L 580501 hipk |
| chr3L | 580932 | 580933 | hipk | 17.9 | 24.1 | 21 | chr3L 580933 hipk |
| **chr3L** | **8970787** | **8970788** | **CG5026** | **17.9** | **16.7** | **17.3** | **chr3L 8970788 CG5026** |
| chr3L | 8993382 | 8993383 | smg | 25.9 | 15.3 | 20.6 | chr3L 8993383 smg |
| chr3L | 9074752 | 9074753 | Tequila | 29.8 | 19.5 | 24.65 | chr3L 9074753 Tequila |
| **chr3L** | **9103274** | **9103275** | **bol** | **23.9** | **26.3** | **25.1** | **chr3L 9103275 bol** |
| chr3L | 9136454 | 9136455 | Use1 | 25 | 16 | 20.5 | chr3L 9136455 Use1 |
| chr3L | 9669496 | 9669497 | fry | 18.2 | 18.2 | 18.2 | chr3L 9669497 fry |
| chr3L | 9945905 | 9945906 | CG34356 | 19.4 | 32.1 | 25.75 | chr3L 9945906 CG34356 |
| chr3R | 10158991 | 10158992 | Invadolysin | 17.9 | 18.2 | 18.05 | chr3R 10158992 Invadolysin |
| chr3R | 10862820 | 10862821 | CG6574 | 20.8 | 33.3 | 27.05 | chr3R 10862821 CG6574 |
| chr3R | 10877257 | 10877258 | CR45195 | 32.7 | 19.5 | 26.1 | chr3R 10877258 CR45195 |
| chr3R | 10889941 | 10889942 | Leash | 18.2 | 18.8 | 18.5 | chr3R 10889942 Leash |
| **chr3R** | **13224637** | **13224638** | **Ace** | **24.8** | **33.3** | **29.05** | **chr3R 13224638 Ace** |
| **chr3R** | **13227870** | **13227871** | **Ace** | **26.6** | **35.3** | **30.95** | **chr3R 13227871 Ace** |
| **chr3R** | **13227970** | **13227971** | **Ace** | **33.42** | **47.95** | **40.69** | **chr3R 13227971 Ace** |
| chr3R | 14330745 | 14330746 | NK7.1 | 16.2 | 16.7 | 16.45 | chr3R 14330746 NK7.1 |
| chr3R | 14660839 | 14660840 | Hexim | 17.1 | 20 | 18.55 | chr3R 14660840 Hexim |
| chr3R | 14669984 | 14669985 | Meltrin | 16.95 | 20.7 | 18.82 | chr3R 14669985 Meltrin |
| chr3R | 14746335 | 14746336 | jvl | 36.4 | 20.9 | 28.65 | chr3R 14746336 jvl |
| chr3R | 14746546 | 14746547 | smp-30 | 45.1 | 24 | 34.55 | chr3R 14746547 smp-30 |
| **chr3R** | **14804349** | **14804350** | **btsz** | **16.5** | **26.5** | **21.5** | **chr3R 14804350 btsz** |
| **chr3R** | **15255687** | **15255688** | **CG42404** | **18.2** | **17.6** | **17.9** | **chr3R 15255688 CG42404** |
| chr3R | 15356475 | 15356476 | Atg4b | 15.4 | 20.7 | 18.05 | chr3R 15356476 Atg4b |
| chr3R | 15414730 | 15414731 | Atx2 | 22.82 | 26.25 | 24.54 | chr3R 15414731 Atx2 |
| chr3R | 15414731 | 15414732 | Atx2 | 19 | 22.6 | 20.8 | chr3R 15414732 Atx2 |
| chr3R | 15849417 | 15849418 | cv-d | 18.8 | 22.2 | 20.5 | chr3R 15849418 cv-d |
| **chr3R** | **16611267** | **16611268** | **NPF** | **22.98** | **23.52** | **23.25** | **chr3R 16611268 NPF** |
| chr3R | 16645743 | 16645744 | CG10324 | 26.1 | 35 | 30.55 | chr3R 16645744 CG10324 |
| chr3R | 17090737 | 17090738 | cal1 | 25 | 15.4 | 20.2 | chr3R 17090738 cal1 |
| **chr3R** | **17731884** | **17731885** | **Lgr1** | **20** | **19** | **19.5** | **chr3R 17731885 Lgr1** |
| chr3R | 17802763 | 17802764 | CG17806 | 20.7 | 18.8 | 19.75 | chr3R 17802764 CG17806 |
| chr3R | 19155252 | 19155253 | CG11779 | 15.6 | 19.1 | 17.35 | chr3R 19155253 CG11779 |
| chr3R | 20783186 | 20783187 | Syp | 21.1 | 17 | 19.05 | chr3R 20783187 Syp |
| chr3R | 20797469 | 20797470 | Syp | 15.3 | 20 | 17.65 | chr3R 20797470 Syp |
| chr3R | 20820785 | 20820786 | CG17271 | 31.2 | 19.75 | 25.48 | chr3R 20820786 CG17271 |
| chr3R | 20862212 | 20862213 | CG3822 | 17 | 26.3 | 21.65 | chr3R 20862213 CG3822 |
| chr3R | 20992668 | 20992669 | Calx | 17.8 | 27.5 | 22.65 | chr3R 20992669 Calx |
| chr3R | 21213891 | 21213892 | SNF4Agamm<br>a | 17.2 | 20 | 18.6 | chr3R_21213892_SNF4Agamm<br>a |
| chr3R | 21354284 | 21354285 | mod(mdg4) | 15 | 19 | 17 | chr3R 21354285 mod(mdg4) |
| chr3R | 21527750 | 21527751 | CG7956 | 30.3 | 26.5 | 28.4 | chr3R 21527751 CG7956 |
| chr3R | 21625731 | 21625732 | E2f | 18 | 23.7 | 20.85 | chr3R 21625732 E2f |
| chr3R | 23261698 | 23261699 | orb | 15.2 | 16.1 | 15.65 | chr3R 23261699 orb |
| chr3R | 23681818 | 23681819 | eIF-3p66 | 20.4 | 21.5 | 20.95 | chr3R_23681819_eIF-3p66 |
| **chr3R** | **23698883** | **23698884** | **prt** | **15.4** | **23.4** | **19.4** | **chr3R_23698884_prt** |
| chr3R | 23723418 | 23723419 | CG10365 | 16.1 | 21.1 | 18.6 | chr3R_23723419_CG10365 |
| chr3R | 23732353 | 23732354 | Rpn9 | 20.4 | 16.7 | 18.55 | chr3R_23732354_Rpn9 |
| chr3R | 24664388 | 24664389 | slo | 23.1 | 30 | 26.55 | chr3R_24664389_slo |
| chr3R | 24802164 | 24802165 | polybromo | 29.2 | 20.98 | 25.09 | chr3R_24802165_polybromo |
| chr3R | 24820485 | 24820486 | Saf-B | 15 | 22 | 18.5 | chr3R_24820486_Saf-B |
| chr3R | 25234647 | 25234648 | CG10420 | 22.2 | 34.5 | 28.35 | chr3R_25234648_CG10420 |
| **chr3R** | **26233920** | **26233921** | **CG12290** | **25.6** | **18.4** | **22** | **chr3R_26233921_CG12290** |
| **chr3R** | **28050111** | **28050112** | **CG34362** | **15.6** | **25.6** | **20.6** | **chr3R_28050112_CG34362** |
| chr3R | 28838531 | 28838532 | Apc | 20.4 | 18.8 | 19.6 | chr3R_28838532_Apc |
| chr3R | 29659085 | 29659086 | Dop1R2 | 15.8 | 39.5 | 27.65 | chr3R_29659086_Dop1R2 |
| **chr3R** | **29674515** | **29674516** | **Bub3** | **27.6** | **17.8** | **22.7** | **chr3R_29674516_Bub3** |
| chr3R | 31457698 | 31457699 | Gprk2 | 23.5 | 24.5 | 24 | chr3R_31457699_Gprk2 |
| chr3R | 31841367 | 31841368 | RhoGAP100F | 17.73 | 19 | 18.37 | chr3R_31841368_RhoGAP100F |
| **chr3R** | **5811503** | **5811504** | **CG11000** | **21.3** | **32.4** | **26.85** | **chr3R_5811504_CG11000** |
| **chr3R** | **5811505** | **5811506** | **CG11000** | **20.8** | **25.7** | **23.25** | **chr3R_5811506_CG11000** |
| chr3R | 7126383 | 7126384 | CG10098 | 18.6 | 19.1 | 18.85 | chr3R_7126384_CG10098 |
| chr3R | 8244435 | 8244436 | CG18749 | 15.4 | 16.1 | 15.75 | chr3R_8244436_CG18749 |
| chr3R | 8244435 | 8244436 | CG33722 | 15.4 | 16.1 | 15.75 | chr3R_8244436_CG33722 |
| chr3R | 9416762 | 9416763 | alpha-Man-II | 30.8 | 32.3 | 31.55 | chr3R_9416763_alpha-Man-II |
| chr3R | 9441505 | 9441506 | ps | 37.92 | 33.52 | 35.72 | chr3R_9441506_ps |
| chr3R | 9471135 | 9471136 | CG16779 | 43.05 | 34.85 | 38.95 | chr3R_9471136_CG16779 |
| chr3R | 9525979 | 9525980 | CG8176 | 16.2 | 24.4 | 20.3 | chr3R_9525980_CG8176 |
| chr3R | 9539464 | 9539465 | mura | 21.75 | 20.05 | 20.9 | chr3R_9539465_mura |
| chr3R | 9794186 | 9794187 | CG8516 | 19.4 | 30.8 | 25.1 | chr3R_9794187_CG8516 |
| chr4 | 478956 | 478957 | Asator | 15.1 | 20 | 17.55 | chr4_478957_Asator |
| chr4 | 532906 | 532907 | zfh2 | 20 | 19.7 | 19.85 | chr4_532907_zfh2 |
| chr4 | 92946 | 92947 | pan | 22.5 | 20.8 | 21.65 | chr4_92947_pan |
| chrX | 10309093 | 10309094 | alpha-Man-I | 24.5 | 29 | 26.75 | chrX_10309094_alpha-Man-I |
| **chrX** | **12331302** | **12331303** | **Ten-a** | **34.8** | **34** | **34.4** | **chrX_12331303_Ten-a** |
| chrX | 16075472 | 16075473 | Tob | 23.68 | 29.32 | 26.5 | chrX_16075473_Tob |
| chrX | 16075887 | 16075888 | Tob | 22.2 | 22 | 22.1 | chrX_16075888_Tob |
| chrX | 16075888 | 16075889 | Tob | 21.6 | 24 | 22.8 | chrX_16075889_Tob |
| **chrX** | **16076959** | **16076960** | **Tob** | **20** | **23.3** | **21.65** | **chrX_16076960_Tob** |
| chrX | 16077193 | 16077194 | Tob | 17.85 | 19.15 | 18.5 | chrX_16077194_Tob |
| chrX | 16089317 | 16089318 | Tob | 15.2 | 16.1 | 15.65 | chrX_16089318_Tob |
| **chrX** | **3321068** | **3321069** | **dnc** | **22.7** | **28.9** | **25.8** | **chrX_3321069_dnc** |
| **chrX** | **3342369** | **3342370** | **dnc** | **17.4** | **22.2** | **19.8** | **chrX_3342370_dnc** |
| **chrX** | **6325433** | **6325434** | **CG15894** | **32** | **20.8** | **26.4** | **chrX_6325434_CG15894** |
| **chrX** | **9172798** | **9172799** | **mei-P26** | **17.3** | **23.1** | **20.2** | **chrX_9172799_mei-P26** |
| chrX | 9179452 | 9179453 | mei-P26 | 17.35 | 22.23 | 19.79 | chrX_9179453_mei-P26 |
| chrX | 9188891 | 9188892 | mei-P26 | 15.8 | 18.4 | 17.1 | chrX_9188892_mei-P26 |

## BIBLIOGRAPHY

Alberti S, Gladfelter A, Mittag T (2019) Considerations and Challenges in Studying Liquid-Liquid Phase Separation and Biomolecular Condensates. Cell 176: 419–434

Alberti S, Mateju D, Mediani L, Carra S (2017) Granulostasis: Protein Quality Control of RNP Granules. Frontiers in Molecular Neuroscience 10

Andrusiak MG, Sharifnia P, Lyu X, Wang Z, Dickey AM, Wu Z, Chisholm AD, Jin Y (2019) Inhibition of Axon Regeneration by Liquid-like TIAR-2 Granules. Neuron 104: 290–304.e298

Ash PEA, Lei S, Shattuck J, Boudeau S, Carlomagno Y, Medalla M, Mashimo BL, Socorro G, Al-Mohanna LFA, Jiang L et al (2021) TIA1 potentiates tau phase separation and promotes generation of toxic oligomeric tau. Proceedings of the National Academy of Sciences 118: e2014188118

Azkanaz M, Rodríguez López A, De Boer B, Huiting W, Angrand P-O, Vellenga E, Kampinga HH, Bergink S, Martens JH, Schuringa JJ et al (2019) Protein quality control in the nucleolus safeguards recovery of epigenetic regulators after heat shock. eLife 8

Babinchak WM, Surewicz WK (2020) Liquid–Liquid Phase Separation and Its Mechanistic Role in Pathological Protein Aggregation. Journal of Molecular Biology 432: 1910–1925

Bah A, Forman-Kay JD (2016) Modulation of Intrinsically Disordered Protein Function by Post-translational Modifications. Journal of Biological Chemistry 291: 6696–6705

Bah A, Vernon RM, Siddiqui Z, Krzeminski M, Muhandiram R, Zhao C, Sonenberg N, Kay LE, Forman-Kay JD (2015) Folding of an intrinsically disordered protein by phosphorylation as a regulatory switch. Nature 519: 106–109

Bakthavachalu B, Huelsmeier J, Sudhakaran IP, Hillebrand J, Singh A, Petrauskas A, Thiagarajan D, Sankaranarayanan M, Mizoue L, Anderson EN et al (2018) RNP-Granule Assembly via Ataxin-2 Disordered Domains Is Required for Long-Term Memory and Neurodegeneration. Neuron 98: 754–766.e754

Becker LA, Huang B, Bieri G, Ma R, Knowles DA, Jafar-Nejad P, Messing J, Kim HJ, Soriano A, Auburger G et al (2017) Therapeutic reduction of ataxin-2 extends lifespan and reduces pathology in TDP-43 mice. Nature 544: 367–371

Berlow RB, Dyson HJ, Wright PE (2015) Functional advantages of dynamic protein disorder. FEBS Lett 589: 2433–2440

Bevilacqua PC, Williams AM, Chou H-L, Assmann SM (2022) RNA multimerization as an organizing force for liquid–liquid phase separation. RNA 28: 16–26

Biogen, 2021. https://clinicaltrials.gov/ct2/show/NCT04494256.

Boeynaems S, Dorone Y, Marian A, Shabardina V, Huang G, Kim G, Sanyal A, Şen N-E, Docampo R, Ruiz-Trillo I, et al (2021) Poly(A)-binding protein is an ataxin-2 chaperone that emulsifies biomolecular condensates. bioRxiv: 2021.2008.2023.457426

Brandmann T, Fakim H, Padamsi Z, Youn JY, Gingras AC, Fabian MR, Jinek M (2018) Molecular architecture of LSM14 interactions involved in the assembly of mRNA silencing complexes. Embo j 37

Buchan JR (2014) mRNP granules. RNA Biology 11: 1019–1030

Calabretta S, Richard S (2015) Emerging Roles of Disordered Sequences in RNA-Binding Proteins. Trends Biochem Sci 40: 662–672

Cao X, Jin X, Liu B (2020) The involvement of stress granules in aging and aging-associated diseases. Aging Cell 19

Chou A, Krukowski K, Jopson T, Zhu PJ, Costa-Mattioli M, Walter P, Rosi S (2017) Inhibition of the integrated stress response reverses cognitive deficits after traumatic brain injury. Proc Natl Acad Sci U S A 114: E6420–e6426

Cirulli ET, Lasseigne BN, Petrovski S, Sapp PC, Dion PA, Leblond CS, Couthouis J, Lu YF, Wang Q, Krueger BJ et al (2015) Exome sequencing in amyotrophic lateral sclerosis identifies risk genes and pathways. Science 347: 1436–1441

Couthouis J, Hart MP, Erion R, King OD, Diaz Z, Nakaya T, Ibrahim F, Kim H-J, Mojsilovic-Petrovic J, Panossian S et al (2012) Evaluating the role of the FUS/TLS-related gene EWSR1 in amyotrophic lateral sclerosis. Human Molecular Genetics 21: 2899–2911

Damrath E, Heck MV, Gispert S, Azizov M, Nowock J, Seifried C, Rüb U, Walter M, Auburger G (2012) ATXN2-CAG42 Sequesters PABPC1 into Insolubility and Induces FBXW8 in Cerebellum of Old Ataxic Knock-In Mice. PLoS Genetics 8: e1002920

De Graeve F, Besse F (2018) Neuronal RNP granules: from physiological to pathological assemblies. Biological Chemistry 399: 623–635

Decker CJ, Teixeira D, Parker R (2007) Edc3p and a glutamine/asparagine-rich domain of Lsm4p function in processing body assembly in Saccharomyces cerevisiae. J Cell Biol 179: 437–449

Deo RC, Bonanno JB, Sonenberg N, Burley SK (1999) Recognition of Polyadenylate RNA by the Poly(A)-Binding Protein. Cell 98: 835–845

Elden AC, Kim HJ, Hart MP, Chen-Plotkin AS, Johnson BS, Fang X, Armakola M, Geser F, Greene R, Lu MM et al (2010) Ataxin-2 intermediate-length polyglutamine expansions are associated with increased risk for ALS. Nature 466: 1069–1075

Formicola N, Vijayakumar J, Besse F (2019) Neuronal ribonucleoprotein granules: Dynamic sensors of localized signals. Traffic 20: 639–649

Gilks N, Kedersha N, Ayodele M, Shen L, Stoecklin G, Dember LM, Anderson P (2004) Stress granule assembly is mediated by prion-like aggregation of TIA-1. Mol Biol Cell 15: 5383–5398

Ginsberg SD, Galvin JE, Chiu TS, Lee VM, Masliah E, Trojanowski JQ (1998) RNA sequestration to pathological lesions of neurodegenerative diseases. Acta Neuropathol 96: 487–494

Gomes E, Shorter J (2019) The molecular language of membraneless organelles. J Biol Chem 294: 7115–7127

Hachet O, Ephrussi A (2004) Splicing of oskar RNA in the nucleus is coupled to its cytoplasmic localization. Nature 428: 959–963

Halliday M, Radford H, Zents KAM, Molloy C, Moreno JA, Verity NC, Smith E, Ortori CA, Barrett DA, Bushell M et al (2017) Repurposed drugs targeting eIF2α-P-mediated translational repression prevent neurodegeneration in mice. Brain 140: 1768–1783

Han TW, Kato M, Xie S, Wu LC, Mirzaei H, Pei J, Chen M, Xie Y, Allen J, Xiao G et al (2012) Cell-free formation of RNA granules: bound RNAs identify features and components of cellular assemblies. Cell 149: 768–779

Harlen KM, Churchman LS (2017) The code and beyond: transcription regulation by the RNA polymerase II carboxy-terminal domain. Nature Reviews Molecular Cell Biology 18: 263–273

Hetz C, Zhang K, Kaufman RJ (2020) Mechanisms, regulation and functions of the unfolded protein response. Nature Reviews Molecular Cell Biology 21: 421–438

Hochberg-Laufer H, Schwed-Gross A, Neugebauer KM, Shav-Tal Y (2019) Uncoupling of nucleo-cytoplasmic RNA export and localization during stress. Nucleic Acids Research 47: 4778–4797

Hofweber M, Dormann D (2019) Friend or foe—Post-translational modifications as regulators of phase separation and RNP granule dynamics. Journal of Biological Chemistry 294: 7137–7150

Huelsmeier J, Walker E, Bakthavachalu B, Ramaswami M (2021) A C-terminal ataxin-2 disordered region promotes Huntingtin protein aggregation and neurodegeneration in Drosophila models of Huntington’s disease. G3 (Bethesda) 11

Hyman AA, Weber CA, Jülicher F (2014) Liquid-Liquid Phase Separation in Biology. Annual Review of Cell and Developmental Biology 30: 39–58

Inagaki H, Hosoda N, Tsuiji H, Hoshino S-I (2020) Direct evidence that Ataxin-2 is a translational activator mediating cytoplasmic polyadenylation. Journal of Biological Chemistry 295: 15810–15825

Ivanov P, Kedersha N, Anderson P (2019) Stress Granules and Processing Bodies in Translational Control. Cold Spring Harb Perspect Biol 11

Jain A, Vale RD (2017) RNA phase transitions in repeat expansion disorders. Nature 546: 243–247

Järvelin AI, Noerenberg M, Davis I, Castello A (2016) The new (dis)order in RNA regulation. Cell Commun Signal 14: 9

Jiang S, Fagman JB, Chen C, Alberti S, Liu B (2020) Protein phase separation and its role in tumorigenesis. eLife 9

Jiménez-López D, Guzmán P (2014) Insights into the evolution and domain structure of ataxin-2 proteins across eukaryotes. BMC Research Notes 7: 453

Kaehler C, Isensee J, Nonhoff U, Terrey M, Hucho T, Lehrach H, Krobitsch S (2012) Ataxin-2-like is a regulator of stress granules and processing bodies. PLoS One 7: e50134

Kato M, Han TW, Xie S, Shi K, Du X, Wu LC, Mirzaei H, Goldsmith EJ, Longgood J, Pei J et al (2012) Cell-free formation of RNA granules: low complexity sequence domains form dynamic fibers within hydrogels. Cell 149: 753–767

Kedersha N, Anderson P (2007) Mammalian stress granules and processing bodies. Methods Enzymol 431: 61–81

Kedersha N, Panas MD, Achorn CA, Lyons S, Tisdale S, Hickman T, Thomas M, Lieberman J, McInerney GM, Ivanov P et al (2016) G3BP-Caprin1-USP10 complexes mediate stress granule condensation and associate with 40S subunits. J Cell Biol 212: 845–860

Kedersha NL, Gupta M, Li W, Miller I, Anderson P (1999) RNA-Binding Proteins Tia-1 and Tiar Link the Phosphorylation of Eif-2α to the Assembly of Mammalian Stress Granules. Journal of Cell Biology 147: 1431–1442

Khong A, Parker R (2020) The landscape of eukaryotic mRNPs. RNA 26: 229–239

Kiebler MA, Bassell GJ (2006) Neuronal RNA granules: movers and makers. Neuron 51: 685–690

Kim G, Gautier O, Tassoni-Tsuchida E, Ma XR, Gitler AD (2020) ALS Genetics: Gains, Losses, and Implications for Future Therapies. Neuron 108: 822–842

Kim H-J, Raphael AR, Ladow ES, Mcgurk L, Weber RA, Trojanowski JQ, Lee VM-Y, Finkbeiner S, Gitler AD, Bonini NM (2014) Therapeutic modulation of eIF2α phosphorylation rescues TDP-43 toxicity in amyotrophic lateral sclerosis disease models. Nature Genetics 46: 152–160

Kim HJ, Kim NC, Wang YD, Scarborough EA, Moore J, Diaz Z, MacLea KS, Freibaum B, Li S, Molliex A et al (2013) Mutations in prion-like domains in hnRNPA2B1 and hnRNPA1 cause multisystem proteinopathy and ALS. Nature 495: 467–473

Kim TH, Payliss BJ, Nosella ML, Lee ITW, Toyama Y, Forman-Kay JD, Kay LE (2021) Interaction hot spots for phase separation revealed by NMR studies of a CAPRIN1 condensed phase. Proc Natl Acad Sci U S A 118

Knowles RB, Sabry JH, Martone ME, Deerinck TJ, Ellisman MH, Bassell GJ, Kosik KS (1996) Translocation of RNA Granules in Living Neurons. The Journal of Neuroscience 16: 7812

Kozlov G, Safaee N, Rosenauer A, Gehring K (2010) Structural basis of binding of P-body-associated proteins GW182 and ataxin-2 by the Mlle domain of poly(A)-binding protein. J Biol Chem 285: 13599–13606

Kwon I, Kato M, Xiang S, Wu L, Theodoropoulos P, Mirzaei H, Han T, Xie S, Corden Jeffry L, McKnight Steven L (2013) Phosphorylation-Regulated Binding of RNA Polymerase II to Fibrous Polymers of Low-Complexity Domains. Cell 155: 1049–1060

Latonen L (2019) Phase-to-Phase With Nucleoli – Stress Responses, Protein Aggregation and Novel Roles of RNA. Frontiers in Cellular Neuroscience 13: 151

Lee J, Yoo E, Lee H, Park K, Hur JH, Lim C (2017) LSM12 and ME31B/DDX6 Define Distinct Modes of Posttranscriptional Regulation by ATAXIN-2 Protein Complex in Drosophila Circadian Pacemaker Neurons. Mol Cell 66: 129–140.e127

Lee KH, Zhang P, Kim HJ, Mitrea DM, Sarkar M, Freibaum BD, Cika J, Coughlin M, Messing J, Molliex A et al (2016) C9orf72 Dipeptide Repeats Impair the Assembly, Dynamics, and Function of Membrane-Less Organelles. Cell 167: 774–788.e717

Li YR, King OD, Shorter J, Gitler AD (2013) Stress granules as crucibles of ALS pathogenesis. J Cell Biol 201: 361–372

Lim C, Allada R (2013) ATAXIN-2 activates PERIOD translation to sustain circadian rhythms in Drosophila. Science 340: 875–879

Lin Y, Currie SL, Rosen MK (2017) Intrinsically disordered sequences enable modulation of protein phase separation through distributed tyrosine motifs. J Biol Chem 292: 19110–19120

Lin Y, Protter SW, David, Rosen K, Michael, Parker R (2015) Formation and Maturation of Phase-Separated Liquid Droplets by RNA-Binding Proteins. Molecular Cell 60: 208–219

Liu EY, Cali CP, Lee EB (2017) RNA metabolism in neurodegenerative disease. Dis Model Mech 10: 509–518

Machida K, Shigeta T, Yamamoto Y, Ito T, Svitkin Y, Sonenberg N, Imataka H (2018) Dynamic interaction of poly(A)-binding protein with the ribosome. Scientific Reports 8: 17435

Mallucci GR, Klenerman D, Rubinsztein DC (2020) Developing Therapies for Neurodegenerative Disorders: Insights from Protein Aggregation and Cellular Stress Responses. Annual Review of Cell and Developmental Biology 36: 165–189

Mandrioli J, Mediani L, Alberti S, Carra S (2020) ALS and FTD: Where RNA metabolism meets protein quality control. Semin Cell Dev Biol 99: 183–192

Mangus DA, Evans MC, Jacobson A (2003) Poly(A)-binding proteins: multifunctional scaffolds for the post-transcriptional control of gene expression. Genome Biol 4: 223

Maniatis T, Reed R (2002) An extensive network of coupling among gene expression machines. Nature 416: 499–506

Martin KC, Ephrussi A (2009) mRNA localization: gene expression in the spatial dimension. Cell 136: 719–730

Matheny T, Van Treeck B, Huynh TN, Parker R (2021) RNA partitioning into stress granules is based on the summation of multiple interactions. RNA 27: 174–189

McCann C, Holohan EE, Das S, Dervan A, Larkin A, Lee JA, Rodrigues V, Parker R, Ramaswami M (2011) The Ataxin-2 protein is required for microRNA function and synapse-specific long-term olfactory habituation. Proc Natl Acad Sci U S A 108: E655–662

McMahon AC, Rahman R, Jin H, Shen JL, Fieldsend A, Luo W, Rosbash M (2016) TRIBE: Hijacking an RNA-Editing Enzyme to Identify Cell-Specific Targets of RNA-Binding Proteins. Cell 165: 742–753

Murray DT, Kato M, Lin Y, Thurber KR, Hung I, McKnight SL, Tycko R (2017) Structure of FUS Protein Fibrils and Its Relevance to Self-Assembly and Phase Separation of Low-Complexity Domains. Cell 171: 615–627.e616

Murthy AC, Dignon GL, Kan Y, Zerze GH, Parekh SH, Mittal J, Fawzi NL (2019) Molecular interactions underlying liquid−liquid phase separation of the FUS low-complexity domain. Nature Structural & Molecular Biology 26: 637–648

Nonhoff U, Ralser M, Welzel F, Piccini I, Balzereit D, Yaspo ML, Lehrach H, Krobitsch S (2007) Ataxin-2 interacts with the DEAD/H-box RNA helicase DDX6 and interferes with P-bodies and stress granules. Mol Biol Cell 18: 1385–1396

Patel A, Lee HO, Jawerth L, Maharana S, Jahnel M, Hein MY, Stoynov S, Mahamid J, Saha S, Franzmann TM et al (2015) A Liquid-to-Solid Phase Transition of the ALS Protein FUS Accelerated by Disease Mutation. Cell 162: 1066–1077

Preissler S, Ron D (2019) Early Events in the Endoplasmic Reticulum Unfolded Protein Response. Cold Spring Harb Perspect Biol 11

Protter DSW, Parker R (2016) Principles and Properties of Stress Granules. Trends Cell Biol 26: 668–679

Protter DSW, Rao BS, Van Treeck B, Lin Y, Mizoue L, Rosen MK, Parker R (2018) Intrinsically Disordered Regions Can Contribute Promiscuous Interactions to RNP Granule Assembly. Cell Rep 22: 1401–1412

Ramaswami M, Taylor JP, Parker R (2013) Altered ribostasis: RNA-protein granules in degenerative disorders. Cell 154: 727–736

Rayman JB, Karl KA, Kandel ER (2018) TIA-1 Self-Multimerization, Phase Separation, and Recruitment into Stress Granules Are Dynamically Regulated by Zn2+. Cell Reports 22: 59–71

Saito M, Hess D, Eglinger J, Fritsch AW, Kreysing M, Weinert BT, Choudhary C, Matthias P (2019) Acetylation of intrinsically disordered regions regulates phase separation. Nat Chem Biol 15: 51–61

Satterfield TF, Pallanck LJ (2006) Ataxin-2 and its Drosophila homolog, ATX2, physically assemble with polyribosomes. Hum Mol Genet 15: 2523–2532

Schuller AP, Wojtynek M, Mankus D, Tatli M, Kronenberg-Tenga R, Regmi SG, Dip PV, Lytton-Jean AKR, Brignole EJ, Dasso M et al (2021) The cellular environment shapes the nuclear pore complex architecture. Nature 598: 667–671

Scoles DR, Meera P, Schneider MD, Paul S, Dansithong W, Figueroa KP, Hung G, Rigo F, Bennett CF, Otis TS et al (2017) Antisense oligonucleotide therapy for spinocerebellar ataxia type 2. Nature 544: 362–366

Shin Y, Brangwynne CP (2017) Liquid phase condensation in cell physiology and disease. Science 357

Shulman JM, Feany MB (2003) Genetic Modifiers of Tauopathy in Drosophila. Genetics 165: 1233–1242

Sidrauski C, Mcgeachy AM, Ingolia NT, Walter P (2015) The small molecule ISRIB reverses the effects of eIF2α phosphorylation on translation and stress granule assembly. eLife 4

Singh A, Hulsmeier J, Kandi AR, Pothapragada SS, Hillebrand J, Petrauskas A, Agrawal K, Rt K, Thiagarajan D, Jayaprakashappa D et al (2021) Antagonistic roles for Ataxin-2 structured and disordered domains in RNP condensation. eLife 10

Strome S, Wood WB (1982) Immunofluorescence visualization of germ-line-specific cytoplasmic granules in embryos, larvae, and adults of Caenorhabditis elegans. Proc Natl Acad Sci U S A 79: 1558–1562

Sudhakaran IP, Hillebrand J, Dervan A, Das S, Holohan EE, Hülsmeier J, Sarov M, Parker R, VijayRaghavan K, Ramaswami M (2014) FMRP and Ataxin-2 function together in long-term olfactory habituation and neuronal translational control. Proc Natl Acad Sci U S A 111: E99–E108

Taylor JP, Brown RH, Cleveland DW (2016) Decoding ALS: from genes to mechanism. Nature 539: 197–206

Tillotson R, Selfridge J, Koerner MV, Gadalla KKE, Guy J, De Sousa D, Hector RD, Cobb SR, Bird A (2017) Radically truncated MeCP2 rescues Rett syndrome-like neurological defects. Nature 550: 398–401

Toretsky JA, Wright PE (2014) Assemblages: functional units formed by cellular phase separation. J Cell Biol 206: 579–588

Trapnell C, Pachter L, Salzberg SL (2009) TopHat: discovering splice junctions with RNA-Seq. Bioinformatics 25: 1105–1111

Van Treeck B, Parker R (2018) Emerging Roles for Intermolecular RNA-RNA Interactions in RNP Assemblies. Cell 174: 791–802

Van Treeck B, Protter DSW, Matheny T, Khong A, Link CD, Parker R (2018) RNA self-assembly contributes to stress granule formation and defining the stress granule transcriptome. Proc Natl Acad Sci U S A 115: 2734–2739

Vogler TO, Wheeler JR, Nguyen ED, Hughes MP, Britson KA, Lester E, Rao B, Betta ND, Whitney ON, Ewachiw TE et al (2018) TDP-43 and RNA form amyloid-like myo-granules in regenerating muscle. Nature 563: 508–513

Wang F, Li J, Fan S, Jin Z, Huang C (2020) Targeting stress granules: A novel therapeutic strategy for human diseases. Pharmacological Research 161: 105143

Wheeler JR, Matheny T, Jain S, Abrisch R, Parker R (2016) Distinct stages in stress granule assembly and disassembly. eLife 5

Wolozin B, Ivanov P (2019) Stress granules and neurodegeneration. Nature Reviews Neuroscience 20: 649–666

Wong YL, Lebon L, Edalji R, Lim HB, Sun C, Sidrauski C (2018) The small molecule ISRIB rescues the stability and activity of Vanishing White Matter Disease eIF2B mutant complexes. eLife 7

Xie J, Kozlov G, Gehring K (2014) The “tale” of poly(A) binding protein: the MLLE domain and PAM2-containing proteins. Biochim Biophys Acta 1839: 1062–1068

Yang P, Mathieu C, Kolaitis RM, Zhang P, Messing J, Yurtsever U, Yang Z, Wu J, Li Y, Pan Q et al (2020) G3BP1 Is a Tunable Switch that Triggers Phase Separation to Assemble Stress Granules. Cell 181: 325–345.e328

Yi H, Park J, Ha M, Lim J, Chang H, Kim VN (2018) PABP Cooperates with the CCR4-NOT Complex to Promote mRNA Deadenylation and Block Precocious Decay. Mol Cell 70: 1081–1088.e1085

Yokoshi M, Li Q, Yamamoto M, Okada H, Suzuki Y, Kawahara Y (2014) Direct binding of Ataxin-2 to distinct elements in 3’ UTRs promotes mRNA stability and protein expression. Mol Cell 55: 186–198

Yoshida M, Yoshida K, Kozlov G, Lim NS, De Crescenzo G, Pang Z, Berlanga JJ, Kahvejian A, Gehring K, Wing SS et al (2006) Poly(A) binding protein (PABP) homeostasis is mediated by the stability of its inhibitor, Paip2. The EMBO Journal 25: 1934–1944

Youn JY, Dyakov BJA, Zhang J, Knight JDR, Vernon RM, Forman-Kay JD, Gingras AC (2019) Properties of Stress Granule and P-Body Proteomes. Mol Cell 76: 286–294

Zhang K, Daigle JG, Cunningham KM, Coyne AN, Ruan K, Grima JC, Bowen KE, Wadhwa H, Yang P, Rigo F et al (2018) Stress Granule Assembly Disrupts Nucleocytoplasmic Transport. Cell 173: 958–971.e917

Zhang Y, Ling J, Yuan C, Dubruille R, Emery P (2013) A role for Drosophila ATX2 in activation of PER translation and circadian behavior. Science 340: 879–882

Zyryanova AF, Kashiwagi K, Rato C, Harding HP, Crespillo-Casado A, Perera LA, Sakamoto A, Nishimoto M, Yonemochi M, Shirouzu M et al (2021) ISRIB Blunts the Integrated Stress Response by Allosterically Antagonising the Inhibitory Effect of Phosphorylated eIF2 on eIF2B. Mol Cell 81: 88–103.e106

